# DNA nanodevice for analysis of force-activated protein extension and interactions

**DOI:** 10.1101/2024.10.25.620262

**Authors:** Kun Zhou, Minhwan Chung, Jing Cheng, John T. Powell, Qi Yan, Jun Liu, Yong Xiong, Martin A. Schwartz, Chenxiang Lin

## Abstract

**Abstract:** Force-induced changes in protein structure and function mediate cellular responses to mechanical stresses. Existing methods to study protein conformation under mechanical force are incompatible with biochemical and structural analysis. Taking advantage of DNA nanotechnology, including the well-defined geometry of DNA origami and programmable mechanics of DNA hairpins, we built a nanodevice to apply controlled forces to proteins. This device was used to study the R1-R2 segment of the talin1 rod domain as a model protein, which comprises two alpha-helical bundles that reversibly unfold under tension to expose binding sites for the cytoskeletal protein vinculin. Electron microscopy confirmed tension-dependent protein extension while biochemical analysis demonstrated enhanced vinculin binding under tension. The device could also be used in pull down assays with cell lysates, which identified filamins as novel tension-dependent talin binders. The DNA nanodevice is thus a valuable addition to the molecular toolbox for studying mechanosensitive proteins.

## Introduction

Forces from muscle and non-muscle cell contraction, gravity, and fluid flow are fundamental determinants of embryonic development, postnatal physiology and many diseases including hypertension, osteoporosis, fibrosis, cancer and atherosclerosis.^1,2^ Cells use a diverse set of proteins to sense and respond to endogenously generated and externally applied mechanical forces. Mechanosensitive proteins are thought to change their conformation under mechanical stress, leading to altered ligand binding, enzymatic activity or ion conductance, eliciting downstream signals that control cell structure, movement and gene expression.^3-5^ Elucidating how forces influence protein structure and function is thus central to understanding mechanically regulated cell behaviors and human physiology and disease.

Existing technologies offer limited means to probe the effects of mechanical loads on protein structure and biochemistry. Fluorescence based live-cell imaging can measure tension across proteins but does not resolve protein structure.^6-10^ Single-molecule techniques such as atomic force microscopy^11^, optical tweezers^12^, and magnetic tweezers^13^ can precisely measure the protein folding/unfolding under defined forces but are largely incompatible with structural and biochemical methods. DNA nanotechnology^14,15^, which generates shape-defined nanostructures from self-assembling DNA strands, offers an approach to fill this need. DNA nanostructures can not only control the spatial arrangement of biomolecules^16-18^, but also mechanically manipulate nucleic acids and proteins^19-21^. They can be produced in large quantities (micro-to milligram scale)^22,23^, enabling a wide selection of analytical methods. Here, we describe a DNA nanodevice to exert tensional stress on a mechanosensitive protein for studying force-regulated protein conformations and molecular interactions.

Integrin-mediated adhesions represent a classic example of a mechanosensitive structure. Integrins initiate formation of dynamic, multi-protein assemblies connecting the intracellular actin network to the extracellular matrix.^24^ Talin is a cytosolic protein that directly connects the integrin beta subunit cytoplasmic domain to F-actin, via its N- and C-terminal domains (NTD and CTD), respectively. Talin’s CTD consists of a string of multi-α helix bundles (R1, R2… R13) that reversibly fold/unfold under tensional forces to conceal/reveal binding sites to an array of ligands. Talin thus not only transmits mechanical force but acts as a mechanochemical switch via opening and closing of these domains.^25^ Remarkably, nine of the rod domains contain buried vinculin binding sites (VBSs) that are exposed upon force-induced domain unfolding. Recruitment of vinculin, which itself contains an F-actin binding site, thus establishes additional attachments to actin, which strengthen the connection between the integrin and the cytoskeleton.

In this study, we used the R1-R2 segment of talin1 as a test subject. These two α helix bundles have a unique side-to-side association to form a single domain that in total contains 3 buried VBSs.^26^ We demonstrate that our DNA nanodevice extends this protein under force and modulates its protein interactions. These data thus demonstrate the utility of this approach for analysis of protein structure and interactions on a biochemical scale.

## Results and Discussion

We designed the protein-tensioning DNA nanodevice with two major components (**Fig. 1**). The first component is a DNA-origami U-shaped frame with a 49-nm wide, 17-nm deep cavity (**Fig. 1a**, caDNAno^27^ design in **Fig. S1**). We initially tested a V-shape origami (inspired by ref^28^) and a rhombus-shape origami (**Fig. S2**) but eventually settled on the U-frame for its superior structural rigidity and homogeneity. Similar U-shape structures have been used as force clamps to control tension across DNA motifs^20,29,30^ or as goniometer for structural determination of DNA-binding proteins^31^. Distinct from the previous work, applying tension to proteins necessitated a second, structure-switching component. To this end, we extended a DNA handle from each side of the cavity, leaving a gap to suspend a protein of interest in the DNA origami frame. A handle can contain two segments of complementary sequences to encode a hairpin structure and function as a “loaded spring”. This handle is kept as extended double-stranded DNA until toehold-mediated strand displacement (TMSD) triggers formation of the hairpin (**Fig. S3**), generating tension on the suspended protein (**Fig. 1a, 1c**). The sequence and length of the spring determine the magnitude of force and the maximal protein extension.

**Figure 1.**
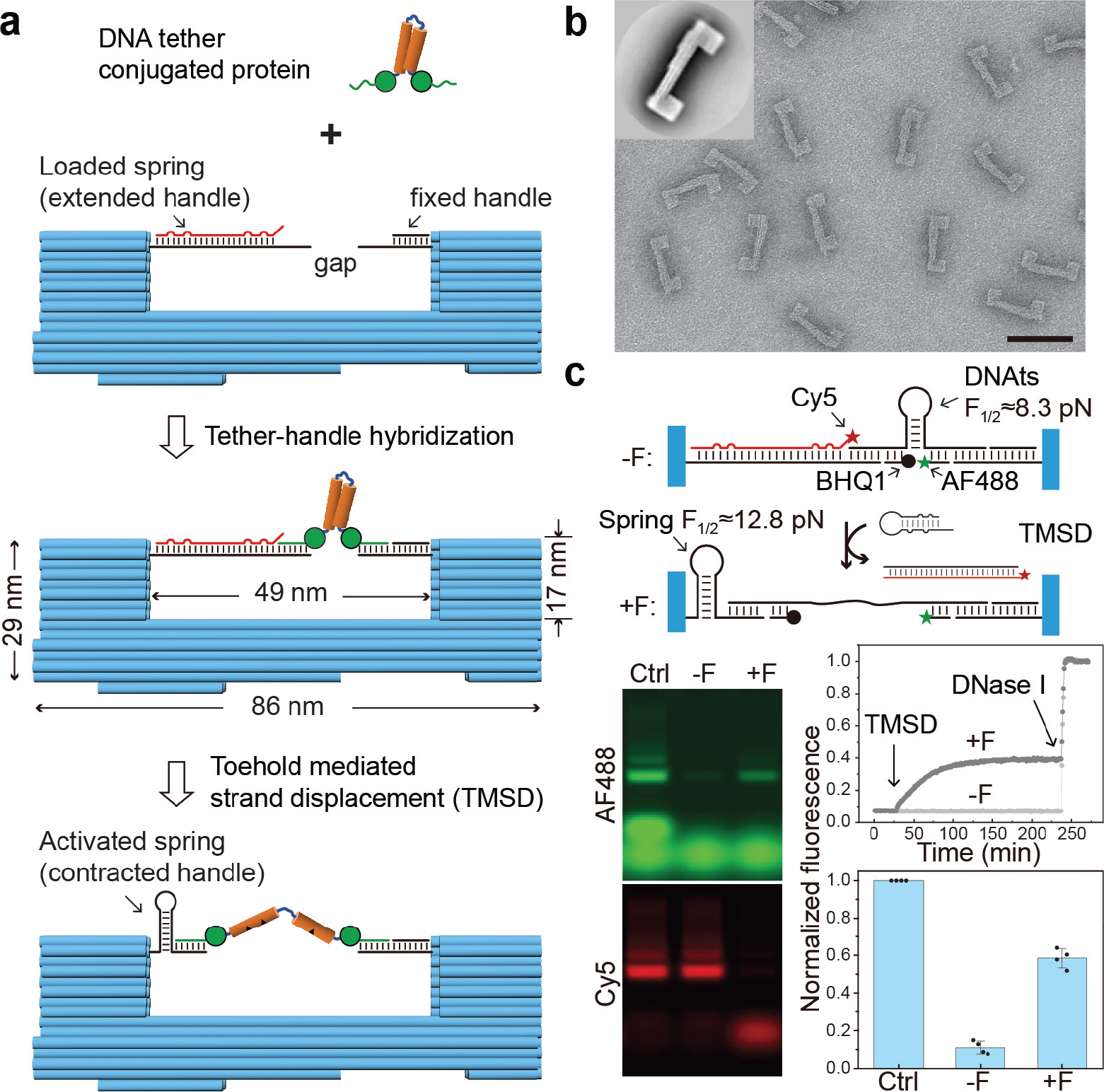
DNA origami device design, assembly and validation. **a)** Schematics of the DNA origami nanomechanical device, which contains a U-shape frame (blue) and a pair of DNA handles with a gap for protein loading. A mechanosensitive protein (e.g., talin R1-R2, orange) can be mounted in the device through site-specifically conjugated DNA tethers (green curls). Controllable force can be generated by transforming an extended DNA handle (loaded spring) into a hairpin (activated spring) via toehold-mediated strand displacement (TMSD) that releases the target strand (red). **b)** A representative negative stain TEM image of the U-frame. Scale bar: 100 nm. Inset: a class average TEM image (120×120 nm^2^). **c)** Validation of force generation using a FRET-based DNA tension sensor (DNAts), a stem-loop structure (predicted unfolding force ≈ 8.3 pN at 25°C) labeled with Alexa Fluor 488 (AF488, green star) and Black Hole Quencher 1 (BHQ1, black circle). Top: schematic showing the TMSD reaction that releases the Cy5 (red star) labeled target strand to activate the DNA spring and unfold the DNAts. Bottom left: images of an agarose gel (upper: AF488 channel, lower: Cy5 channel) in which the DNA devices loaded with DNAts missing BHQ1 (Ctrl) and with complete DNAts before (-F) and 3 hr after (+F) TMSD were electrophoresed. Bottom right: trace of DNAts fluorescence during application of tension (upper, normalized to nuclease-treated DNAts), and the DNAts fluorescence quantified from the gel image (lower, normalized to DNAts without BHQ1). The bar graph shows mean and standard deviation.

Following standard DNA origami folding and purification workflow (see Methods)^32,33^, we obtained the DNA origami frame with the designed geometry (**Fig. 1b**). To test its ability to generate force, we incorporated a tension sensor—a DNA stem-loop labeled with a fluorophore (Alexa Flour 488)-quencher (Blackhole Quencher 1) pair (similar to ref^10^)—into the U-frame in tandem with an extended spring (**Fig. 1c**). We left a 9-nm initial gap between DNA handles to accommodate the sensor (see a coarse-grained model by oxDNA^34^ in **Fig. S4**) and designed a DNA spring that contracts by 14 nm upon formation of a hairpin with a theoretical unfolding force of ∼12.8 pN (F_1/2_, defined as the force under which half of the hairpins unfold^35^). Displacing the Cy5-labeled target strand activated the spring and unzipped the DNA tension sensor (DNAts). Monitoring fluorescence of the DNAts showed that the system reached equilibrium within ∼2 hours. Gel electrophoresis confirmed that the TMSD is nearly complete after 3 hours and that the fluorescence of DNAts increased to ∼50% of the unquenched level (i.e., device loaded with quencher-free DNAts, **Fig. 1c**). The incomplete recovery is likely the combined result of steeper transition energy barrier near the unfolded state of a hairpin that leads to the weaker spring folding force than its theoretical F_1/2_,^35,36^ considerable tension (∼6 pN based on a worm-like chain model^37^) on the unfolded DNAts that prompts refolding, and some misassembled DNA devices failing to properly exert force (**Fig. S5**). Nevertheless, this result demonstrates that the nanodevice generates tension sufficient to unfold half of the sensor (F_1/2_), which is predicted at 8.3 pN. As expected, the same DNA origami device unzipped a DNAts with smaller F_1/2_ (6.1 pN) with higher efficiency but only minimally unfolded a DNAts with F_1/2_ comparable to the spring (**Fig. S6**). Considering the average tension across talin in cells is ∼6 pN,^38^ with peaks above 7–11 pN,^39^ we expect the DNA nanodevice to unfold talin R1-R2 and activate vinculin binding.

To incorporate talin R1-R2 into the DNA device with defined orientation, we expressed R1-R2 with a SNAP-tag and a HaloTag at its N- and C-termini, respectively (S-R1-R2-H). We then reacted it with a benzylguanine modified 22 nucleotide (nt) tether and a cholorohexane modified 24 nt tether to generate doubly labeled S-R1-R2-H (**Fig. 2a**). The purified conjugation product showed clear band shifts upon binding with oligonucleotides complementary to the tethers (C1, C2, or both, see **Fig. 2b**), confirming over 90% yield of doubly labeled S-R1-R2-H as we recently reported^40^. DNA-guided self-assembly of the components (U-frame, DNA spring, and dual-labeled protein) yielded R1-R2 containing DNA nanodevices (Origami-R1-R2). After removing free proteins and DNA origami aggregates by rate-zonal centrifugation^41^ (**Fig. 2c**), we imaged the purified DNA nanodevices by negative-stain transmission electron microscopy (TEM). Most devices (74.1%) contained a slightly off-centered protein in the cavity as expected, while a small portion (2.4%) appeared protein-free and another 23.5% contained erroneously captured proteins (e.g., one at each end of the cavity, **Fig. 2d**). Occasionally visible DNA handles on both sides of S-R1-R2-H further confirmed its proper suspension in the nanodevice (as shown in some zoomed-in micrographs in **Fig. 2d** and **Fig. 3**).

**Figure 2.**
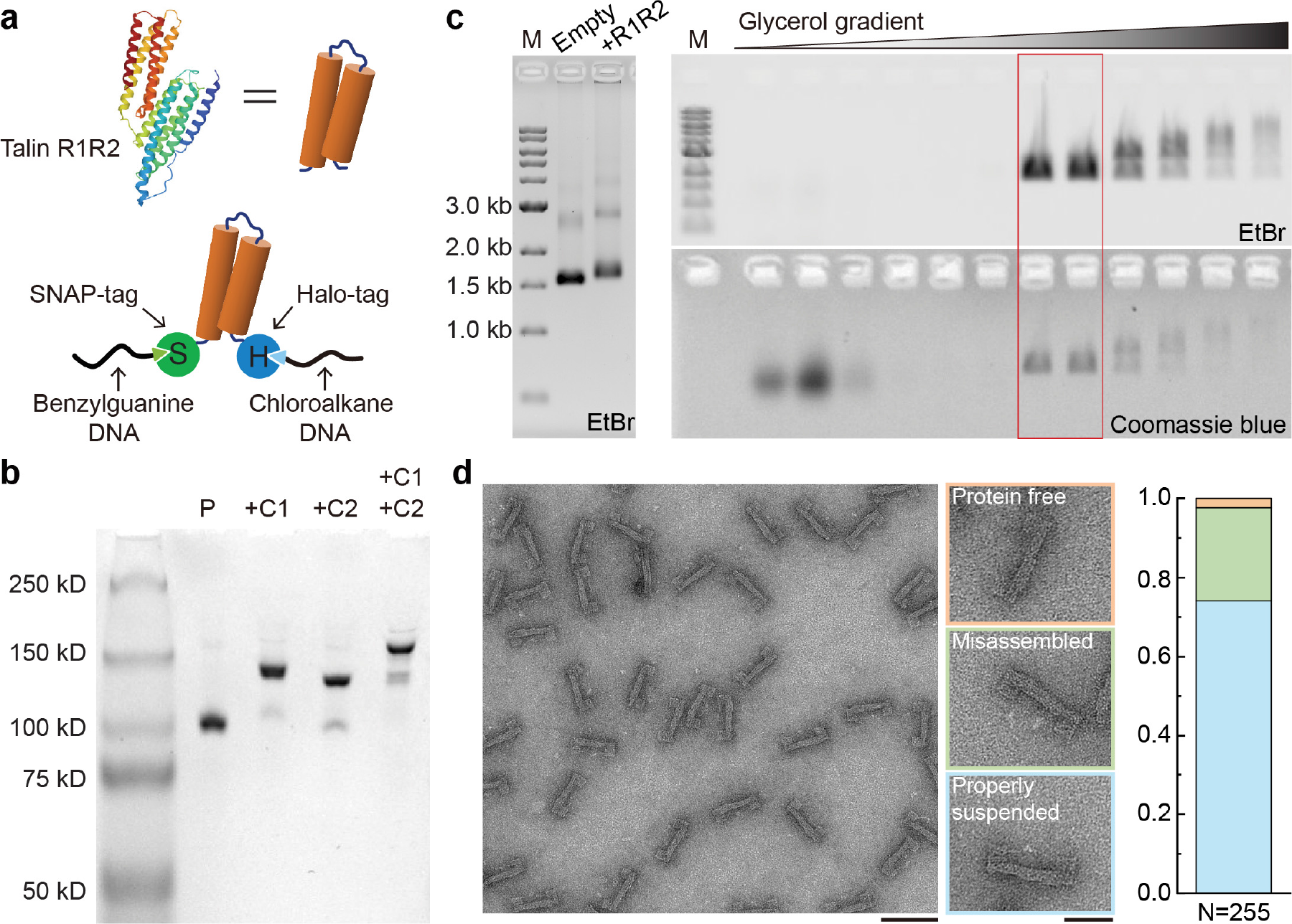
Assembly of protein-loaded DNA-origami device. **a)** Cartoon of the talin R1-R2 domain (PDB 1SJ8, top) and schematic of R1-R2 with terminal SNAP- and Halo-Tags (S-R1-R2-H) conjugated to benzylguanine- and chloroalkane-labeled DNA tethers (bottom). **b)** Native PAGE showing purified S-R1-R2-H (P), and DNA conjugated S-R1-R2-H hybridized to a 100-nt DNA strand complementary to the benzylguanine tether (+C1), to an 80-nt DNA strand complementary to the chloroalkane tether (+C2), or to both complementary strands (+C1 +C2). **c)** Left: agarose gel resolving the DNA origami device before (empty) and after (+R1R2) protein loading; right: agarose gel resolving fractions recovered from a glycerol gradient after rate-zonal centrifugation. Fractions in the red box are enriched in protein-loaded devices. M: 1kb DNA ladder; EtBr: ethidium bromide stain. **d)** Negative-stain electron micrographs of purified DNA-origami device after S-R1-R2-H loading. Among 255 devices analyzed, 2.4% are without a protein (orange group), 23.5% erroneously captured proteins (green group), and 74.1% contain a properly suspended protein (blue group). Scale bars: 100 nm for the zoomed-out image and 50 nm for zoomed-in images.

**Figure 3.**
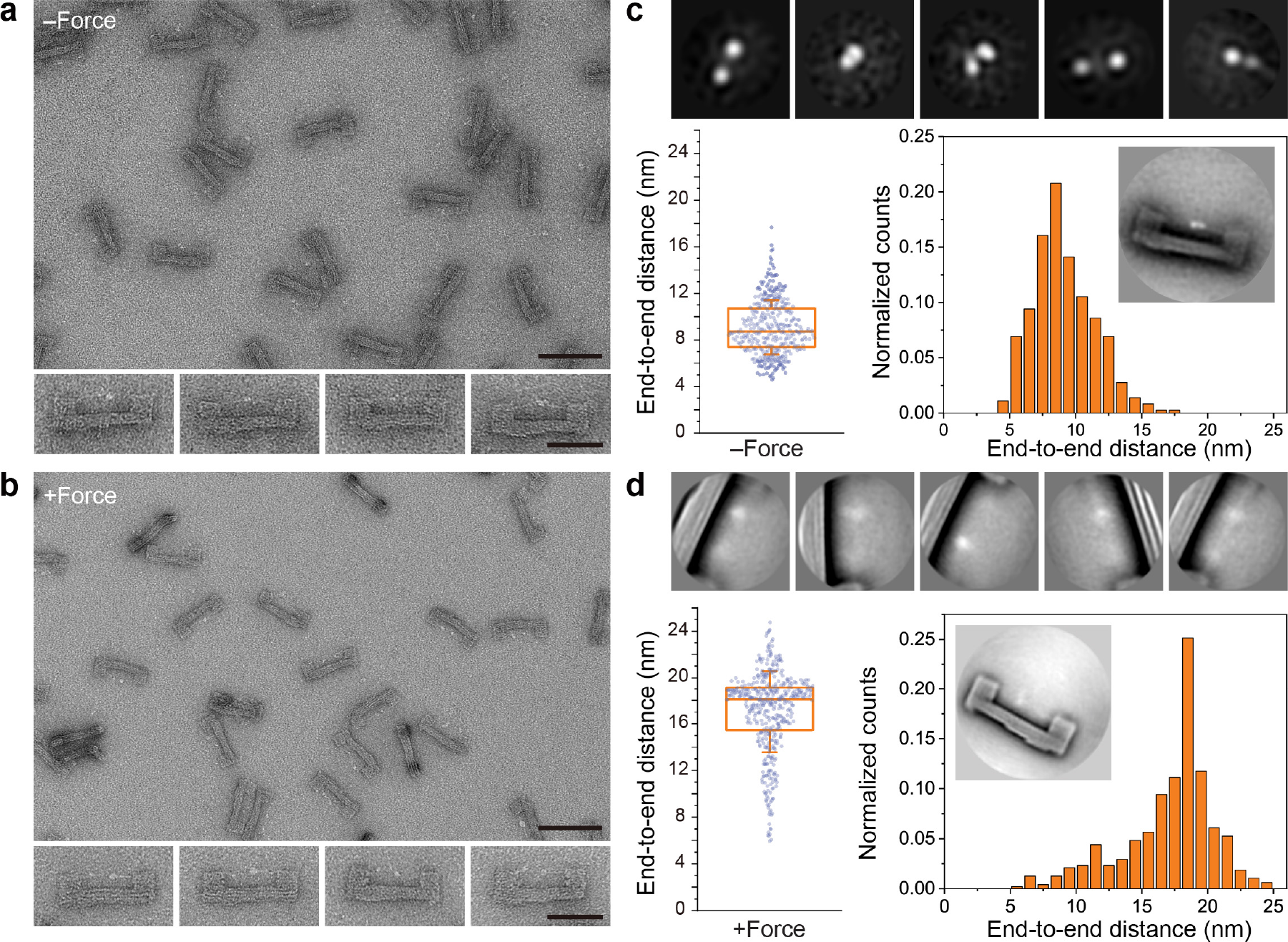
Tension induced conformational changes of S-R1-R2-H. **a)** Negative-stain TEM images of purified DNA origami devices containing S-R1-R2-H. The initial gap for protein loading is 6 nm. The theoretical unfolding force of DNA spring is ∼12.8 pN. Scale bars: 100 nm for zoomed-out image and 50 nm for zoomed-in images. **b)** Same as **a**, but after TMSD triggered force application. **c)** Single-particle analyses of S-R1-R2-H in the relaxed state. Top row: gallery of 2D class averages (30×30 nm^2^). Bottom row: distribution of distances measured between the two farthest domains of protein suspended within a DNA origami device (n = 361). The box and whisker plot shows the median, 25% and 75% percentiles, and standard deviation. Inset shows a class average (120×120 nm^2^) of the DNA device with relaxed S-R1-R2-H. **d)** Same as **c**, but after force induced protein unfolding. 2D class averages in the top row are 50×50 nm^2^. For the distance distribution, the number of devices measured is 483.

Under negative-stain TEM, S-R1-R2-H in the DNA-origami device appeared as two or three closely placed puncta (**Fig. 3a** and **S7**). The different appearances are probably because of internal movement among the protein domains (i.e., HaloTag, SNAP-tag and R1-R2) and the rotation of the entire protein relative to the DNA frame. With resolution limited by negative staining technique, we measured 9.1±2.3 nm (n=361) end-to-end distance (*d*) of relaxed S-R1-R2-H (**Fig. 3b** and **S7**), in agreement with an AlphaFold model^42^ and TEM images of free S-R1-R2-H (**Fig. S8**). After force application most of the S-R1-R2-H molecules appeared as two asymmetric, well-separated puncta (*d*=17.1±3.5 nm, n=483) (**Fig. 3c, 3d**). We attribute the two puncta to HaloTag (33 kDa) and SNAP-tag (19 kDa), respectively, suggesting tension-induced unfolding of the R1-R2 domains (33 kDa). The larger distribution in *d* compared to the relaxed state reflects conformational fluctuations under tension, likely due to comparable unfolding energy of R1-R2 and folding energy of the DNA spring. The average change in *d* (*Δd:* ∼8 nm) is smaller than the designed spring contraction length (14 nm), suggesting slack in the initial protein handle assembly may have buffered the protein extension. We note that the unfolding force of talin R1-R2 was measured to be 15–25 pN using magnetic tweezers^43^, exceeding the folding force of the DNA spring (< 13 pN). However, unlike the magnetic tweezers that apply constantly increasing force in short bursts (<10 seconds), the DNA nanodevice subjected the protein to mechanical load for as long as the device remains intact (hours). It is well known that the force required for protein unfolding is proportional to logarithm of the loading rate^44,45^. Thus, our results are consistent with established principles.

A distinct advantage of the DNA nanodevices over conventional mechanical manipulation tools is its compatibility with bulk biochemical methods. We set out to test the tension-dependent molecular interaction of R1-R2 using purified vinculin head domain (VinD1) as a binding partner. For this purpose, we incubated Origami-R1-R2 with purified VinD1 labeled with FLAG-tag, pulled down biotin-labeled DNA devices by streptavidin-coated magnetic beads, washed and analyzed the proteins retained on the beads (**Fig. 4a**). We first varied the gap between the DNA handles from 2–16 nm while maintaining the DNA spring unfolding force (∼12.8 pN) and length (48 nt). In addition, we built a defective device with R1-R2 attached to only one handle (“1-handle” in **Fig. 4b, 4c**) to serve as a no force control. Western blots show that VinD1 binding was negligible for R1-R2 on nanodevices before force application (marked as -F groups in **Fig. 4b**) as well as on the no force device, confirming that loading R1-R2 into DNA device did not disrupt its folding state. Upon TMSD and hairpin formation (+F groups in **Fig. 4b**), VinD1 binding increased up to 6-fold for R1-R2 mounted within an initial gap of 6 or 9 nm (but not 2 or 16 nm). This is because long handles that leave a very small gap attenuated force transmission to R1-R2 while shorter handles are too far apart for anchorage of a relaxed S-R1-R2-H at both ends. Keeping the initial gap at 6 nm, we then varied the design of the spring to change the force it generates. We tested three handles of equal lengths (48 nt) but with different sequences. Two of them encode hairpin structures with theoretical unfolding forces of 12.8 pN and 8.8 pN (**Fig. 4c**, strong and week HP, respectively), and the third encodes no secondary structure with a maximal tension of ∼6.1 pN (**Fig. 4c**, unstructured). Both hairpin-based springs led to increased VinD1 binding upon activation, in contrast to the unstructured handle, which did not trigger binding. Therefore, our data show that tensional force in the range of 6.1–8.8 pN is sufficient to expose VBSs in R1-R2 when the protein is under sustained mechanical load. This range is likely an overestimation, as flexible components such as the peptide linkers between R1-R2 and the conjugation tags may dampen the force. Comparing the biochemistry data and the structural analyses (**Fig. 3**) further suggests that a force-induced R1-R2 extension by ∼8 nm is coupled to significant increase in vinculin binding.

**Figure 4.**
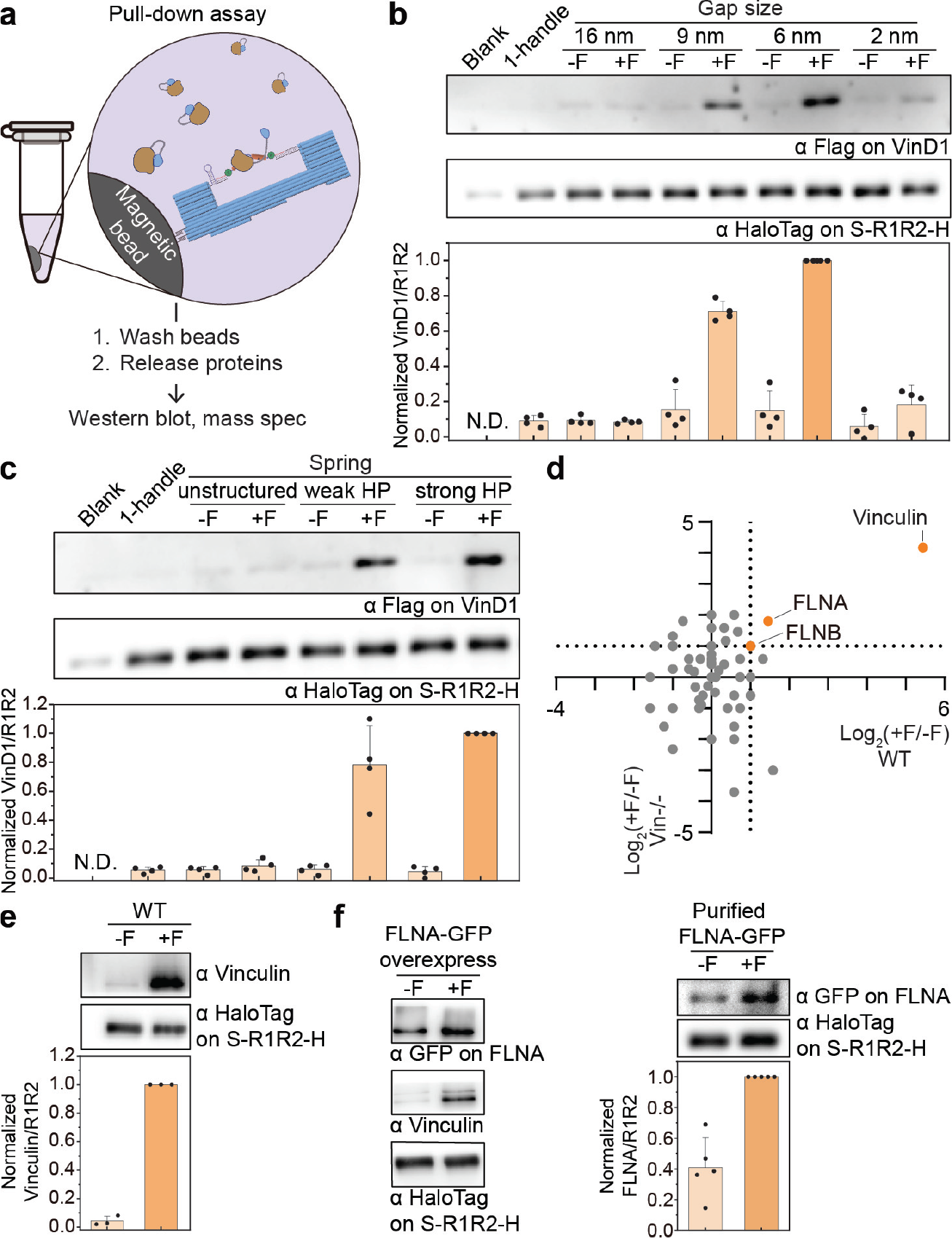
Force-dependent protein interactions with talin R1-R2. **a)** Schematic of the pulldown assay. DNA origami devices, loaded with R1-R2, are immobilized on magnetic beads to assay force-activated protein binding. **b)** Western blot showing force-activated binding of VinD1-FLAG to R1-R2 as a function of gap size. A DNA hairpin with unfolding force (F_1/2_) of ∼12.8 pN serves as spring. **c)** Western blot showing the effect of DNA spring design on force-activated VinD1-FLAG binding to R1-R2. An unstructured spring and two hairpin (HP) forming springs (F_1/2_≈ 8.8 and 12.8 pN), all 48-nt long, were tested. Initial gap = 6 nm. **d)** Tension-dependent R1-R2 binding proteins in cell lysates detected by mass spectrometry. Ratios of R1-R2 binders in the presence (+F) and absence (-F) of tension are plotted for the wildtype (WT, x-axis) and vinculin depleted (Vin-/-, y-axis) NIH3T3 cells. **e)** Western blot showing force-activated R1-R2 binding to vinculin in lysates of WT NIH3T3 cells. **f)** Western blots validating force-activated R1-R2 binding to FLNA in lysates of FLNA-GFP overexpressing cells (left) and purified FLNA-GFP (right). The ratios of talin binders to R1-R2 are normalized to the 6 nm gap, 12.8 pN HP, +F condition for each trial and plotted as bar graphs, which show means and standard deviations. Blank: DNA device without handle; 1-handle: defective device with R1-R2 tethered to only one handle; N.D.: not detected.

Having established its ability to mechanically induce protein extension and interactions, we asked whether the nanodevice could be used to identify new tension-dependent talin interactors. We mixed bead-immobilized DNA nanodevices containing relaxed or tensioned R1-R2 with NIH3T3 cell lysate and characterized the pulled down cellular proteins using mass spectroscopy-based proteomics (MS). This assay was carried out using lysates from unperturbed cell and vinculin knockdown (∼87% efficiency, **Fig. S9**) cells and results plotted so that proteins in the upper right quadrant represent those that show increased binding in both conditions, thus are likely vinculin independent. Among dozens of pulled-down proteins (**Fig. 4d**), vinculin showed the greatest tension-dependent binding (∼44-fold detected by MS and ∼20-fold by western blot, **Fig 4f**); even the residual vinculin in the knockdown cells bound to force-activated R1-R2. The Filamins (FlnA > B) also showed a tension-dependent increase in binding to talin R1-R2. Filamins are actin crosslinkers known for its role in actin network remodeling.^46^ Our previous work shows that FlnA knockdown in uniaxially stretched cells resulted in reduced actin connectivity and polarized tension distribution on talin.^47^ However, direct FlnA binding to talin has not been reported. Incubation of Origami-R1-R2 with purified FlnA recapitulated the increased association under tension (**Fig. 4f**), demonstrating a direct interaction independent of vinculin.

Quantitative analysis of SDS-PAGE suggests that despite having 3 VBSs, an S-R1-R2-H extended by ∼8 nm bound to ∼0.4 VinD1 on average (**Fig. S10**). This prompted us to reengineer the DNA device for further protein extension. In this design, an S-R1-R2-H is initially suspended by two DNA handles with orthogonal hairpin-encoding sequences (i.e., loaded springs), enabling independent or simultaneous spring activation for programmable protein extension (**Fig. 5a**). Specifically, the left (L) and right (R) springs (**Fig. S3** and **S11**) are designed to contract by 14 and 8 nm upon TMSD-triggered folding, respectively, for a maximal contraction length of 22 nm. Resolving the positions of the conjugation tags (SNAP and Halo) on R1-R2 relative to the asymmetric U-frame confirmed the protein loading and elongation (**Fig. 5b** and **Fig S12**). Activating the L, R, and both springs produced mean *Δd* of 6.8 nm, 3.5 nm, and 10.1 nm, respectively, in comparison with the relaxed S-R1-R2-H (*d*=9.2±2.4 nm). We expected that compared to the single-spring device (**Fig. 3**), activating two springs (F_1/2_≈12.8 and 14.0 pN) in tandem do not exert greater force and the probability for both DNA hairpins to properly fold is lower. This is reflected by the large distribution in *d* when the protein was stretched by both springs (L+/R+ in **Fig. 5b**). Nevertheless, the two springs’ effect on *Δd* were additive, allowing R1-R2 to unfold further to potentially reveal additional VBSs. Indeed, binding assay done on cell lysates showed >2-fold higher vinculin pulldown by R1-R2 stretched by both springs than by either one (**Fig. 5c**). The dual-spring device thus unambiguously correlates R1-R2 conformation with its vinculin binding capability. We recently reported the programmable extension of R1-R2 by a double-stranded DNA (dsDNA) ‘clamp’^40^, where we found significantly increased VinD1 binding only with *Δd* of ∼15 nm (force ≤2.2 pN). In contrast, the hairpins used in this work generate higher force to extend R1-R2. Correspondingly, vinculin binding increased markedly with mean *Δd* as low as 3.5 nm, demonstrating that both stress and strain modulate R1-R2’s protein interactions.

**Figure 5.**
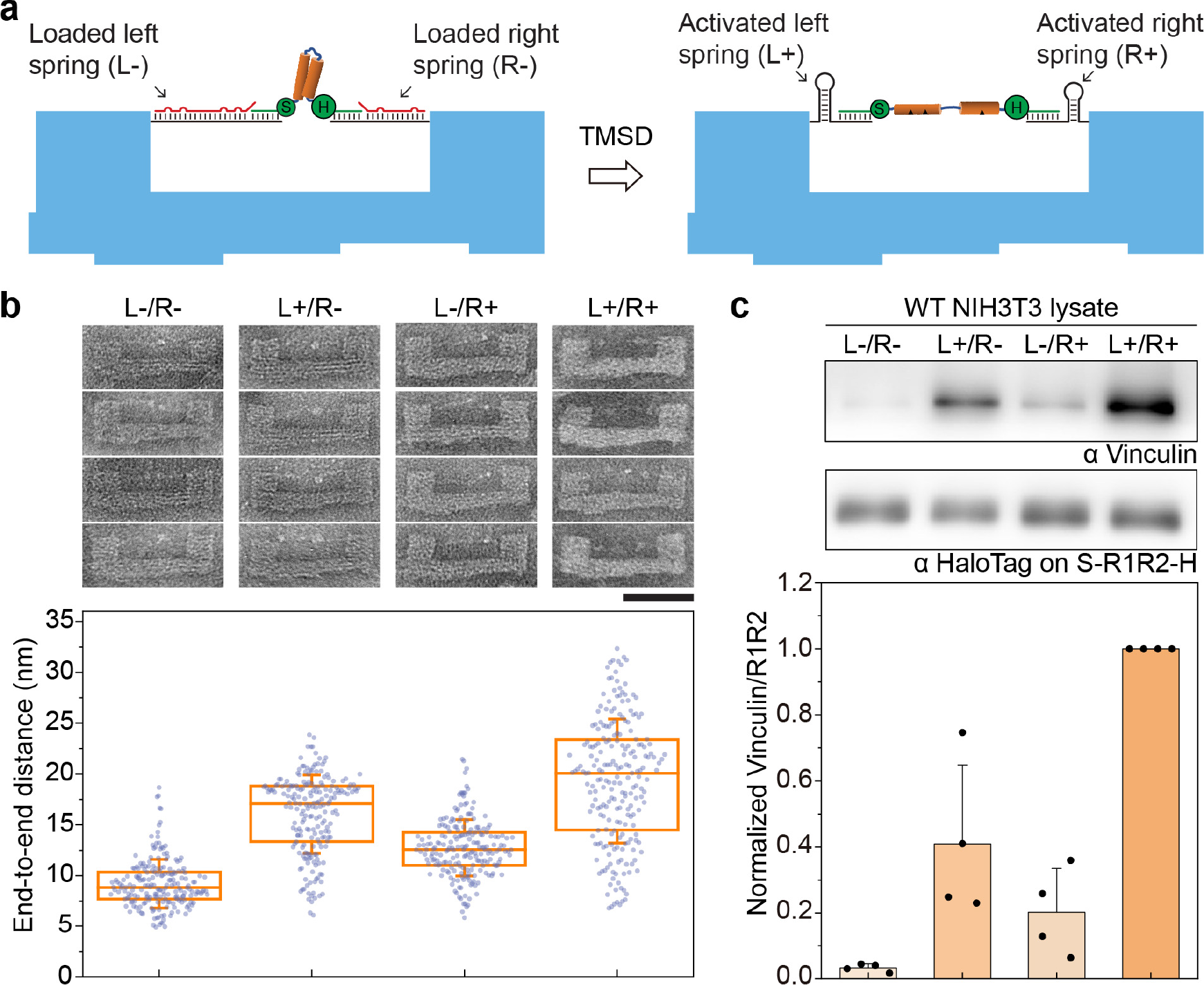
Programmable extension of talin R1-R2 led to varying vinculin binding. **a)** Schematics showing an S-R1-R2-H extended by two springs (L and R) with different lengths and sequences on a DNA origami device. The R1-R2 domain, target strands, protein-conjugated DNA tethers, and DNA origami frame are depicted in orange, red, green, and blue, respectively. **b)** Representative TEM micrographs (top) and measured end-to-end distances (bottom) of S-R1-R2-H under relaxed (L-/R-) and various extended conditions (L+/R-, L-/R+, L+/R+) as the result of selective activation of the springs. The box and whisker plot shows the median, 25% and 75% percentiles, and standard deviation. Number of devices measured are 205, 208, 210 and 220. Scale bar: 50 nm. **c)** Western blot characterization of vinculin binding by relaxed and extended R1-R2 in cell lysates. The ratio of vinculin to R1-R2 is normalized to the maximally extended (L+/R+) condition for each trial and plotted as bar graphs, which show means and standard deviations.

## Conclusions

Motivated by the need for nanomechanical tools compatible with biochemical analyses, we built a DNA origami-based device to apply tension to a protein of interest and analyze effects on its structure and protein interactions. Compared to previous DNA origami structures intended to study protein interactions with tensioned DNA^20,29,30,48^, this device directly exerts force on the protein rather than its nucleic acid substrate. As a proof-of-concept demonstration, folded talin R1-R2 domains were mechanically switched to elongated conformations by a DNA spring, which induced vinculin binding. Interesting, our results show that the strain required for vinculin binding is rather modest (mean *Δd* ≲10 nm), well below the length of a fully extended string of 9 alpha-helices in R1-R2 (>30 nm), suggesting a partial and likely transient unfolding is sufficient to expose VBSs. Using a dual-spring setup, we show higher strain increases vinculin binding. In an unbiased screen for tension-dependent R1-R2 interactors in cell lysate, tensioned R1-R2 showed greater binding to filamins, which also occurred with purified proteins, thus identifying a new tension-modulated talin interaction. Previous work identified a functional link between talin and FlnA in cell responses to applied strains;^47^ whether the interaction identified here mediates or contributes to that effect is an important question for future work.

The protein tensioning nanodevice provides a platform for studying the structure-function relationship of mechanosensitive proteins. Thanks to its modular design, the device can be adapted to study other proteins of interest, for example by changing the width of DNA origami frame and the sequence of DNA spring. Using two site-specifically conjugated DNA strands to guide the protein loading onto the DNA nanodevice, this method is compatible with a broad spectrum of proteins. The different protein-binding behaviors by R1-R2 loaded in devices with various gap size and DNA spring strength highlight the need to optimize the design parameters based on the dimensions and force-response characteristics of the target protein. An important future goal is to obtain a high-resolution protein structure by cryo-EM. However, doing so for talin R1-R2 is limited by its relatively small size and the apparent heterogeneity of the deformation products. Proteins with larger mass and better-defined tension-induced conformations are more suitable for structural determination by TEM. This methodology, in its current or adapted form, is ready for application to a wide range of proteins implicated in cell and tissue responses to mechanical forces.

## Materials and Methods

### DNA origami design and preparation

All DNA oligonucleotides were purchased from Integrated DNA Technologies (IDT). Unmodified staple strands were purchased in a 96-well plate format with concentrations normalized to 100 μM and used without additional purification. Fluorophore-modified strands were purchased with HPLC purification in tube format. Oligonucleotides used to form DNA handles, including hairpin/unstructured springs, target strands, and displacement strands (sequences in **Table S1**), and those longer than 80 nt were purchased in tube format and purified by denaturing polyacrylamide gel electrophoresis (PAGE). The p8064 scaffold strand, an M13mp18-derived single-stranded DNA, was prepared as previously described^32^.

The DNA origami frame was designed using caDNAno^27^. DNA structures were prepared by mixing p8064 scaffold (20 nM) with the corresponding staple strands (each 140 nM) in a folding buffer (5 mM Tris-HCl, 1 mM EDTA, 5 mM NaCl, 16 mM MgCl_2_, pH 8.0). The mixture was then placed in a thermocycler (Bio-Rad) and incubated at 80°C for 5 min, followed by annealing from 65°C to 25 °C in 40 hours.

The assembled structures were purified by PEG precipitation^33^. Briefly, the post-annealing mixture of scaffold and staple strands was mixed with a 16% (w/v) PEG-8000 solution in 40 mM Tris-Base, 500 mM NaCl, 40 mM MgCl_2_, pH 8.0 at a 3:1 volume ratio, held at room temperature (RT) for 15 min, and centrifuged at 15,000g for 15 min at RT. The supernatant was carefully discarded, and the pellet was resuspended in a 1× TE-Mg^2+^ buffer (10 mM Tris-HCl, 1 mM EDTA, 12.5 mM MgCl_2_, pH 8.0). The resuspended structures were PEG precipitated again and dissolved in the 1× TE-Mg^2+^ buffer. The dissolved structures were incubated at 37°C for 30 min and centrifuged at 20,000g for 5 min. The supernatant was collected, and the DNA origami concentration was determined using a NanoDrop 2000 spectrometer (Thermo Fisher Scientific).

### Plasmids

The gene encoding *Mus musculus* talin1 rod domain R1-R2 (amino acid 482–786) was cloned between a N-terminal SNAP-tag gene (Addgene, Plasmid #101135) and a C-terminal HaloTag gene (Promega, G8031) followed by a 6×His-tag sequence. This construct was inserted into the pcDNA3.1 vector using Gibson assembly. The vinculin head D1 domain (amino acid 1–257, VinD1) was cloned into a pcDNA3.1 vector with C-terminal Flag and 6×His-tags. The plasmid for filamin A-GFP (pcDNA3-FLNA-GFP) is a kind gift from Dr. D.A. Calderwood. The plasmid pOPINE2-GFP nanobody was obtained from Addgene (Plasmid #49172).

### Protein expression and purification

SNAP-R1-R2-Halo (S-R1-R2-H), VinD1, and FLNA-GFP plasmids were transfected into 293TX cells cultured in DMEM supplemented with 10% FBS and 1% Penicillin-Streptomycin, using Opti-MEM (Thermo Fisher Scientific) and Lipofectamine 2000 (Invitrogen), following the manufacturer’s protocol. GFP-nanobody was expressed in BL21 chemically competent E. coli (Invitrogen, C600003) by heat-shock at 37°C for 30 seconds. Cells were suspended in 1× PBS, pH 7.4, supplemented with 15 mM imidazole, 1 mM dithiothreitol (DTT), 0.2% Tween-20, protease inhibitor cocktail (Roche, 1183617001). For E. coli expression, 1 mg/mL lysozyme was added to the same buffer (used exclusively for the GFP nanobody). After sonication, the lysate was centrifuged at 30,000g for 30 minutes at 4°C. The supernatant was then incubated with Ni-NTA beads (Qiagen) for 3 hours with gentle agitation at 4°C. The beads were thoroughly washed with the same buffer, and proteins were eluted using 500 mM imidazole in 1× PBS at 4°C. To remove imidazole, we used 7K MWCO Zeba Spin desalting columns (Thermo Scientific), pre-equilibrated with 1× PBS containing 0.05% Tween-20 and 1 mM DTT. The purified proteins were either used immediately for oligonucleotide conjugation (S-R1-R2-H DNA dual-labeling) or aliquoted and snap-frozen for later use. For FLNA-GFP purification, the FLNA-GFP cell lysate was incubated with purified 6×His-tagged GFP nanobody for 2 hours, followed by Ni-NTA bead purification as described above.

### Protein-DNA conjugation

Amino-modified DNA oligonucleotides (sequences in **Table S1**) purchased from IDT were purified by ethanol precipitation and resuspended in Milli-Q ultrapure water at 2 mM concentration. The BG-GLA-NHS (New England Biolabs, S9151S) and HaloTag Succinimidyl Ester (O4) Ligand (PROMEGA, P6751) were reconstituted in DMSO or DMF to a final concentration of 20 mM. The oligonucleotides (32 μL of 2 mM) and labeling ligands (96 μL of 20 mM) were mixed at 1:30 molar ratio in 67 mM HEPES, pH 8.5 buffer, and incubated at RT for 1 hour. Unreacted ligands were removed by ethanol precipitation. The resulting benzylguanine (BG)-DNA and cholorohexane (CL)-DNA were resuspended in Milli-Q water, (optionally) purified using a 7K MWCO Zeba Spin desalting column (Thermo Fisher Scientific) and stored at -20°C until use.

A mixture of purified S-R1-R2-H (10 μM) and the BG-DNA and CL-DNA (30 μM each) in 1× PBS buffer containing 1 mM DTT was incubated at 15°C for 1.5 hours and then at 4°C overnight. Unconjugated oligonucleotides were removed through Ni-NTA bead purification (Qiagen) or size exclusion chromatography using a Superdex 200 10/300 column (GE Healthcare). DNA-tether conjugated proteins were aliquoted and stored at -80°C. DNA dual-labeling efficiency was verified through SDS-PAGE or native PAGE, both stained with Coomassie blue (Thermo Fisher Scientific). For SDS-PAGE, precast NuPAGE Bis-Tris gels (4–12% gradient, Invitrogen) were run in MES-SDS or MOPS-SDS buffer (Invitrogen). For native PAGE, tether-conjugated protein (0.3 μg/μL) was incubated with five-fold excess of a 100-nt DNA strand (C1, partially complementary to BG-tether) and/or an 80-nt DNA strand (C2, partially complementary to CL-tether) at RT for 1 hour before being electrophoresed in a 6% polyacrylamide gel in 1× TBE buffer (89 mM tris base, 89 mM boric acid, 2 mM EDTA, pH 8.3) supplemented with 2 mM MgCl_2_.

### Applying force to S-R1-R2-H by DNA origami device

Purified DNA origami devices (typically 23 nM) was mixed with 3.5 to 5-fold molar excess of dual-DNA labeled S-R1-R2-H and incubated at 37°C for 1 to 1.5 hours. The resulting Origami-R1-R2 assemblies were purified by rate-zonal centrifugation in a 15–45% glycerol gradient^41^. Briefly, 100 μL of Origami-R1-R2 was placed on top of a quasi-linear glycerol gradient (15–45%, in 1× TE-Mg^2+^ buffer) in a 0.8 mL Open-Top Thinwall Ultra-Clear tube (Beckman Coulter) and spun in an SW 55 Ti rotor (Beckman Coulter) at 48,000 RPM at 4°C for 55 min, followed by fraction collection. Fractions (4 μL each) were then loaded onto a 1% agarose gel containing ethidium bromide (EtBr) and run in 0.5× TBE buffer supplemented with 10 mM MgCl_2_. The agarose gel was imaged on a Typhoon FLA 9500 scanner (Cytiva) for the EtBr fluorescence, stained by Coomassie blue, and imaged again. Fractions containing monomeric Origami-R1-R2 were buffer exchanged into 1× TE-Mg^2+^ buffer using an Amicon Ultra centrifugal filter with 100 kD NMWL (Millipore). To trigger TMSD and extend R1-R2, a displacement strand was added to purified Orgaimi-R1-R2 at a 5–7.5 to 1 molar ratio. The mixture was first incubated at 37°C for 1 hour and then at 16°C or RT overnight. Aliquots of samples before and after TMSD were subjected to negative-stain TEM. Unpurified Origami-R1-R2 before and after TMSD was used for pulldown experiments.

### FRET-based DNAts

A DNAts consists of a 120-nt stem-loop forming strand, an AF488-labeled strand and a BHQ1-labeled strand. To form DNAts, the PAGE-purified 120-mer oligonucleotide (2 μM) was heated to 80°C for 10 min and quenched on ice before mixed with the AF488-strand and BHQ1-strand at 1:2:2 molar ratio in a TE buffer (10 mM Tris-HCl, pH 8) supplemented with 100 mM NaCl and 2 mM MgCl_2_; the 3-strand mixture was incubated at 25°C for 2 hours. Unpurified DNAts was added to the DNA-origami device (typically 23 nM) containing a Cy5-labeled target strand at a 5:1 molar ratio unless otherwise specified, and the mixture was incubated at 37°C for 1 to 1.5 hours. To apply force, the DNAts-loaded devices (Origami-DNAts) were mixed with a 7.5-fold molar excess of displacement strand and incubated at 25°C for 3 hr. An ‘always-on’ DNAts was formed using an unmodified oligonucleotide with the same sequence as the BHQ1-strand and loaded into the DNA origami device to serve as a control. The Origami-DNAts samples, before and after force application, were electrophoresed on an agarose gel along with the control sample and imaged for AF488 and Cy5 fluorescence using a Typhoon FLA 9500 scanner. The gel was subsequently stained with EtBr overnight and imaged again.

Real-time monitoring of DNAts fluorescence was performed on a Bio-Tek Synergy HT microplate reader. Briefly, 100 μL of 23 nM Origami-DNAts was purified by PEG precipitation to remove excess DNAts and its components; the pelleted Origami-DNAts was dissolved in 100 μL of 1× TE-Mg^2+^ buffer. The AF488 fluorescence signal of the purified samples (50 μL each, typically 17–20 nM) was measured every minute (excitation: 488 nm, emission: 525 nm) in a 384-well black glass-bottomed microplate (Porvair). To apply force, 5 μL of 1.3–1.5 μM displacement strand was added at the 15–30 min time point. After several hours, 1 μL of 0.2–0.4 U/μL DNAse I (Thermo Fisher Scientific) was added to all samples to dissociate AF488 from BHQ1.

To quantify the unzipped DNAts in DNA origami devices, the fluorescence signals contributed by misassembled Origami-DNAts were subtracted from the total fluorescence detected. Because ∼16% of the DNA devices were estimated to contain two DNAts (**Fig. S5**), the incorrectly incorporated DNAts constituted 2×16%/(1+16%)≈28% of the fluorescence detected from the Origami-DNAts before force application (-F) and the control samples (‘always-on’ and DNase I treated DNAts). Because these misassembled DNAts are not under mechanical load after spring activation, they contribute the same amount of fluorescence before (-F) and after (+F) force application.

### Western blot

Samples were incubated at 98°C for 5 min in 1× Laemmli sample buffer. Proteins were resolved using SDS-PAGE and transferred to a nitrocellulose membrane via wet electroblotting (Bio-Rad). The membrane was blocked with 5% milk (Americanbio, AB10109) in TBS containing 0.1% Tween-20 (TBS-T) and incubated at 4°C overnight with the following primary antibodies diluted in TBS-T: mouse anti-HaloTag (1:8000; Promega, G9211), mouse anti-Flag M2 (1:2000; Millipore Sigma, F1804), anti-vinculin (Proteintech, 26520-1-AP), anti-GFP (Santa Cruz Biotechnology, sc9996), and Rabbit anti-GAPDH (1:4000; Cell signaling, #2118). Membranes were washed three times with TBS-T for 5 minutes each on a shaker at RT and then incubated with HRP-conjugated secondary antibody (anti-mouse (PI-2000-1) or anti-rabbit (PI-1000-1), 1:8000 in TBS-T; Vector Laboratories) for one hour at RT, followed by three more washes with TBS-T. Bands were visualized using chemiluminescence substrates (SuperSignal West Pico Plus or Femto, Thermo Fisher Scientific) and imaged with the G:box system (Syngene).

### Pulldown assay

Typically, FLNA-GFP or VinD1 was first diluted in 100 μL of 1× PBS with 0.2% Tween-20 and 10 mM MgCl_2_, PBS-TM), and then mixed with 100 μL of biotinylated Origami-R1-R2 (23 nM, in 1× TE-Mg^2+^) at a binder:R1-R2 molar ratio of 6–10:1. The mixture was supplemented with 2% BSA and incubated for 90 min at 4°C with mild agitation. Then, 40 μL of pre-washed streptavidin magnetic beads (New England Biolabs, S1420S) were added to the mixture and incubated for an additional 40 min. The beads were gently washed three times with PBS-TM and subsequently treated with DNase I (final concentration 0.02–0.04 U/μL, Thermo Fisher Scientific or New England Biolabs) in DNase I reaction buffer (New England Biolabs, B0303S) containing 0.1% Tween-20 at 37°C for 30 min while shaking. The samples were then analyzed by western blot as described above.

### Vinculin knockdown

Vinculin knockdown NIH3T3 cells were generated using CRISPR-Cas9 with a guide RNA (sgVinculin 5’-GCCGTCAGCAACCTCGTCC-3’), following established procedures^49^. Briefly, lentiviruses for Cas9 and guide RNA delivery were produced by transfecting 293TX cells with packaging vectors (pCMV-VSV-G, #8454; psPAX2, #12260; Addgene) and the pLenti-CRISPR DNA vector (#52961, Addgene) using Lipofectamine 2000 (Invitrogen). Virus-containing supernatant was collected at 36- and 60-hours post-transfection and used to infect NIH3T3 cells for knockout, along with 8 μM polybrene (Sigma). Sixteen hours post-infection, the medium was replaced with fresh medium containing 7.5 μM puromycin for selection, with medium changes every 48 hours. After six days, the selected cells were ready for further experiments.

### Cell lysate pulldown and mass spectrometry

Wildtype and vinculin-deficient NIH3T3 cells or FLN-GFP overexpressing 293TX cells were collected and incubated in ice-cold hypotonic buffer (0.1× PBS, 1× protease inhibitor (Thermo Fisher Scientific, 78430), and 0.05% Tween-20) for 15 min to enrich the cytosolic components in the lysate. The lysates were centrifuged at 21,000 g for 30 min at 4°C, and the supernatants were collected for the pulldown assay. For each condition, 30 μL of the lysate was diluted in 500 μL of PBS-TM containing 1× protease inhibitor and aurintricarboxylic acid (250 μg/mL, Thermo Fisher Scientific, AC103060250). The mixture was pre-cleared with 40 μL of poly-T magnetic beads (NEB, S1419S) for 30 min at 4°C with agitation. After removing the beads, 125 μL of Origami-R1-R2 (23 nM) was added to the lysate, incubated for 40 min at 4°C with mild agitation, pulled-down, and analyzed by western blot as described above. Following DNase I treatment, Laemmli sample buffer was added to the eluate, and the mixture was heated to 95°C for 5 minutes. The samples were loaded onto a 4–20% SDS gradient gel (Bio-Rad, #4561094) and electrophoresed. When the samples had migrated approximately 1 cm into the gel, the run was stopped. The gel was fixed and visualized using Coomassie blue. After destaining, the gel plug was excised and submitted to the Taplin Mass Spectrometry Facility at Harvard Medical School for LC/MS/MS analysis to identify interacting partners.

### Electron microscopy

Typically, 10 μL of 2.5 nM sample solution was adsorbed to a glow-discharged, formvar/carbon-coated copper grid, and blotted away after 2 min using filter paper. The grid was rinsed with TE buffer and then stained with 2% uranyl formate. Images were acquired on a JEOL JEM-1400 Plus microscope operated at 80 kV with a bottom-mount 4k×3k charge-coupled device (CCD) camera (Advanced Microscopy Technologies). The protein end-to-end distance was measured between the centers of the two farthest protein particles using ImageJ software. Particle picking was automated using TOPAZ^50^ unless otherwise specified. Two-dimensional class averages of the picked particles were generated using Relion 4.0.

## Acknowledgements

This work was supported by a National Institutes of Health (NIH) Grant (R01-HL155543) to M.A.S. and C.L., a Department of Defense/Army Research Office MURI Grant (W911NF1410403) to M.A.S., a Smith Family Foundation Odyssey Award to C.L., an NIH grant R01-AI162260 to Y.X. and C.L., NIH grants (R01AI087946, R01AI152421, R01AI132818) to J.L., and a fellowship from the American Heart Association (20POST35080107) to M.C.

## Author Contributions

M.A.S. conceived the project. K.Z., M.C., J.T.P., M.A.S. and C.L. designed the approach. K.Z., M.C., J.T.P. and Q.Y. performed experiments. K.Z., M.C. and J.C. analyzed data. J.L., Y.X., C.L. and M.A.S. provided resources and supervised the study. K.Z. and C.L. wrote the original draft. K.Z., M.C., M.A.S. and C.L. edited the manuscript. All authors participated in the discussions and approved the manuscript. K.Z. and M.C. contributed equally.

## Competing Interests

The authors declare no competing interests.

## Notes

### Competing Interest Statement

The authors have declared no competing interest.

### Summary of Updates

Figure 5 added. Supplementary Materials updated. Other minor changes to the manuscript for clarity and consistency.

## References

1 Mierke, C. T. Extracellular Matrix Cues Regulate Mechanosensing and Mechanotransduction of Cancer Cells. Cells 13 (2024). 10.3390/cells13010096

2 Orr, A. W., Helmke, B. P., Blackman, B. R. & Schwartz, M. A. Mechanisms of mechanotransduction. Dev Cell 10, 11–20 (2006). 10.1016/j.devcel.2005.12.006

3 Hoffman, B. D., Grashoff, C. & Schwartz, M. A. Dynamic molecular processes mediate cellular mechanotransduction. Nature 475, 316–323 (2011). 10.1038/nature10316

4 Jin, P., Jan, L. Y. & Jan, Y. N. Mechanosensitive Ion Channels: Structural Features Relevant to Mechanotransduction Mechanisms. Annu Rev Neurosci 43, 207–229 (2020). 10.1146/annurev-neuro-070918-050509

5 Romani, P., Valcarcel-Jimenez, L., Frezza, C. & Dupont, S. Crosstalk between mechanotransduction and metabolism. Nat Rev Mol Cell Biol 22, 22–38 (2021). 10.1038/s41580-020-00306-w

6 Grashoff, C. et al. Measuring mechanical tension across vinculin reveals regulation of focal adhesion dynamics. Nature 466, 263–266 (2010). 10.1038/nature09198

7 Hu, Y. et al. DNA-based ForceChrono probes for deciphering single-molecule force dynamics in living cells. Cell 187, 3445–3459e3415 (2024). 10.1016/j.cell.2024.05.008

8 Ren, Y. et al. Force redistribution in clathrin-mediated endocytosis revealed by coiled-coil force sensors. Sci Adv 9, eadi1535 (2023). 10.1126/sciadv.adi1535

9 Tao, A. et al. Identifying constitutive and context-specific molecular-tension-sensitive protein recruitment within focal adhesions. Dev Cell 58, 522–534 e527 (2023). 10.1016/j.devcel.2023.02.015

10 Zhang, Y., Ge, C., Zhu, C. & Salaita, K. DNA-based digital tension probes reveal integrin forces during early cell adhesion. Nat Commun 5, 5167 (2014). 10.1038/ncomms6167

11 Fisher, T. E., Oberhauser, A. F., Carrion-Vazquez, M., Marszalek, P. E. & Fernandez, J. M. The study of protein mechanics with the atomic force microscope. Trends Biochem Sci 24, 379–384 (1999). 10.1016/s0968-0004(99)01453-x

12 Bustamante, C. J., Chemla, Y. R., Liu, S. & Wang, M. D. Optical tweezers in single-molecule biophysics. Nat Rev Methods Primers 1 (2021). 10.1038/s43586-021-00021-6

13 Choi, H. K., Kim, H. G., Shon, M. J. & Yoon, T. Y. High-Resolution Single-Molecule Magnetic Tweezers. Annu Rev Biochem 91, 33–59 (2022). 10.1146/annurev-biochem-032620-104637

14 Dey, S. et al. DNA origami. Nat Rev Method Prime 1 (2021). 10.1038/s43586-020-00009-8

15 Seeman, N. C. & Sleiman, H. F. DNA nanotechnology. Nat Rev Mater 3 (2018). 10.1038/natrevmats.2017.68

16 Fisher, P. D. E. et al. A Programmable DNA Origami Platform for Organizing Intrinsically Disordered Nucleoporins within Nanopore Confinement. ACS Nano 12, 1508–1518 (2018). 10.1021/acsnano.7b08044

17 Fu, J. et al. Multi-enzyme complexes on DNA scaffolds capable of substrate channelling with an artificial swinging arm. Nat Nanotechnol 9, 531–536 (2014). 10.1038/nnano.2014.100

18 Zeng, Y. C. et al. Fine tuning of CpG spatial distribution with DNA origami for improved cancer vaccination. Nat Nanotechnol 19, 1055–1065 (2024). 10.1038/s41565-024-01615-3

19 Mills, A. et al. A modular spring-loaded actuator for mechanical activation of membrane proteins. Nat Commun 13, 3182 (2022). 10.1038/s41467-022-30745-2

20 Nickels, P. C. et al. Molecular force spectroscopy with a DNA origami-based nanoscopic force clamp. Science 354, 305–307 (2016). 10.1126/science.aah5974

21 Wang, Y. et al. A nanoscale DNA force spectrometer capable of applying tension and compression on biomolecules. Nucleic Acids Res 49, 8987–8999 (2021). 10.1093/nar/gkab656

22 Praetorius, F. et al. Biotechnological mass production of DNA origami. Nature 552, 84–87 (2017). 10.1038/nature24650

23 Jia, Y. L., Chen, L. M., Liu, J., Li, W. & Gu, H. Z. DNA-catalyzed efficient production of single-stranded DNA nanostructures. Chem-Us 7, 959–981 (2021). 10.1016/j.chempr.2020.12.001

24 Sun, Z., Guo, S. S. & Fassler, R. Integrin-mediated mechanotransduction. J Cell Biol 215, 445–456 (2016). 10.1083/jcb.201609037

25 Goult, B. T., Yan, J. & Schwartz, M. A. Talin as a mechanosensitive signaling hub. J Cell Biol 217, 3776–3784 (2018). 10.1083/jcb.201808061

26 Papagrigoriou, E. et al. Activation of a vinculin-binding site in the talin rod involves rearrangement of a five-helix bundle. EMBO J 23, 2942–2951 (2004). 10.1038/sj.emboj.7600285

27 Douglas, S. M. et al. Rapid prototyping of 3D DNA-origami shapes with caDNAno. Nucleic Acids Res 37, 5001–5006 (2009). 10.1093/nar/gkp436

28 Funke, J. J. & Dietz, H. Placing molecules with Bohr radius resolution using DNA origami. Nat Nanotechnol 11, 47–52 (2016). 10.1038/nnano.2015.240

29 Kramm, K. et al. DNA origami-based single-molecule force spectroscopy elucidates RNA Polymerase III pre-initiation complex stability. Nat Commun 11, 2828 (2020). 10.1038/s41467-020-16702-x

30 Xiong, Q. et al. DNA Origami Post-Processing by CRISPR-Cas12a. Angew Chem Int Ed Engl 59, 3956–3960 (2020). 10.1002/anie.201915555

31 Aksel, T., Yu, Z., Cheng, Y. & Douglas, S. M. Molecular goniometers for single-particle cryo-electron microscopy of DNA-binding proteins. Nat Biotechnol 39, 378–386 (2021). 10.1038/s41587-020-0716-8

32 Douglas, S. M. et al. Self-assembly of DNA into nanoscale three-dimensional shapes. Nature 459, 414–418 (2009). 10.1038/nature08016

33 Wagenbauer, K. F. et al. How We Make DNA Origami. Chembiochem 18, 1873–1885 (2017). 10.1002/cbic.201700377

34 Poppleton, E. et al. Design, optimization and analysis of large DNA and RNA nanostructures through interactive visualization, editing and molecular simulation. Nucleic Acids Res 48, e72 (2020). 10.1093/nar/gkaa417

35 Woodside, M. T. et al. Nanomechanical measurements of the sequence-dependent folding landscapes of single nucleic acid hairpins. Proc Natl Acad Sci U S A 103, 6190–6195 (2006). 10.1073/pnas.0511048103

36 Bercy, M. & Bockelmann, U. Hairpins under tension: RNA versus DNA. Nucleic Acids Res 43, 9928–9936 (2015). 10.1093/nar/gkv860

37 Marko, J. F. & Siggia, E. D. Stretching DNA. Macromolecules 28, 8759–8770 (1995). 10.1021/ma00130a008

38 Kumar, A. et al. Talin tension sensor reveals novel features of focal adhesion force transmission and mechanosensitivity. J Cell Biol 213, 371–383 (2016). 10.1083/jcb.201510012

39 Austen, K. et al. Extracellular rigidity sensing by talin isoform-specific mechanical linkages. Nat Cell Biol 17, 1597–1606 (2015). 10.1038/ncb3268

40 Chung, M., Zhou, K., Powell, J. T., Lin, C. & Schwartz, M. A. DNA-Based Molecular Clamp for Probing Protein Interactions and Structure under Force. ACS Nano 18, 27590–27596 (2024). 10.1021/acsnano.4c08663

41 Lin, C., Perrault, S. D., Kwak, M., Graf, F. & Shih, W. M. Purification of DNA-origami nanostructures by rate-zonal centrifugation. Nucleic Acids Res 41, e40 (2013). 10.1093/nar/gks1070

42 Abramson, J. et al. Accurate structure prediction of biomolecular interactions with AlphaFold 3. Nature 630, 493–500 (2024). 10.1038/s41586-024-07487-w

43 Yao, M. et al. The mechanical response of talin. Nat Commun 7, 11966 (2016). 10.1038/ncomms11966

44 Carrion-Vazquez, M. et al. Mechanical and chemical unfolding of a single protein: a comparison. Proc Natl Acad Sci U S A 96, 3694–3699 (1999). 10.1073/pnas.96.7.3694

45 Evans, E. & Ritchie, K. Strength of a weak bond connecting flexible polymer chains. Biophys J 76, 2439–2447 (1999). 10.1016/S0006-3495(99)77399-6

46 Zhou, J., Kang, X., An, H., Lv, Y. & Liu, X. The function and pathogenic mechanism of filamin A. Gene 784, 145575 (2021). 10.1016/j.gene.2021.145575

47 Kumar, A. et al. Filamin A mediates isotropic distribution of applied force across the actin network. J Cell Biol 218, 2481–2491 (2019). 10.1083/jcb.201901086

48 Aissaoui, N. et al. Modular Imaging Scaffold for Single-Particle Electron Microscopy. ACS Nano 15, 4186–4196 (2021). 10.1021/acsnano.0c05113

49 Driscoll, T. P., Ahn, S. J., Huang, B., Kumar, A. & Schwartz, M. A. Actin flow-dependent and - independent force transmission through integrins. Proc Natl Acad Sci U S A 117, 32413–32422 (2020). 10.1073/pnas.2010292117

50 Bepler, T. et al. Positive-unlabeled convolutional neural networks for particle picking in cryo-electron micrographs. Nat Methods 16, 1153–1160 (2019). 10.1038/s41592-019-0575-8

