## Supplementary Materials for "DNA nanodevice for analysis of force-activated protein extension and interactions"

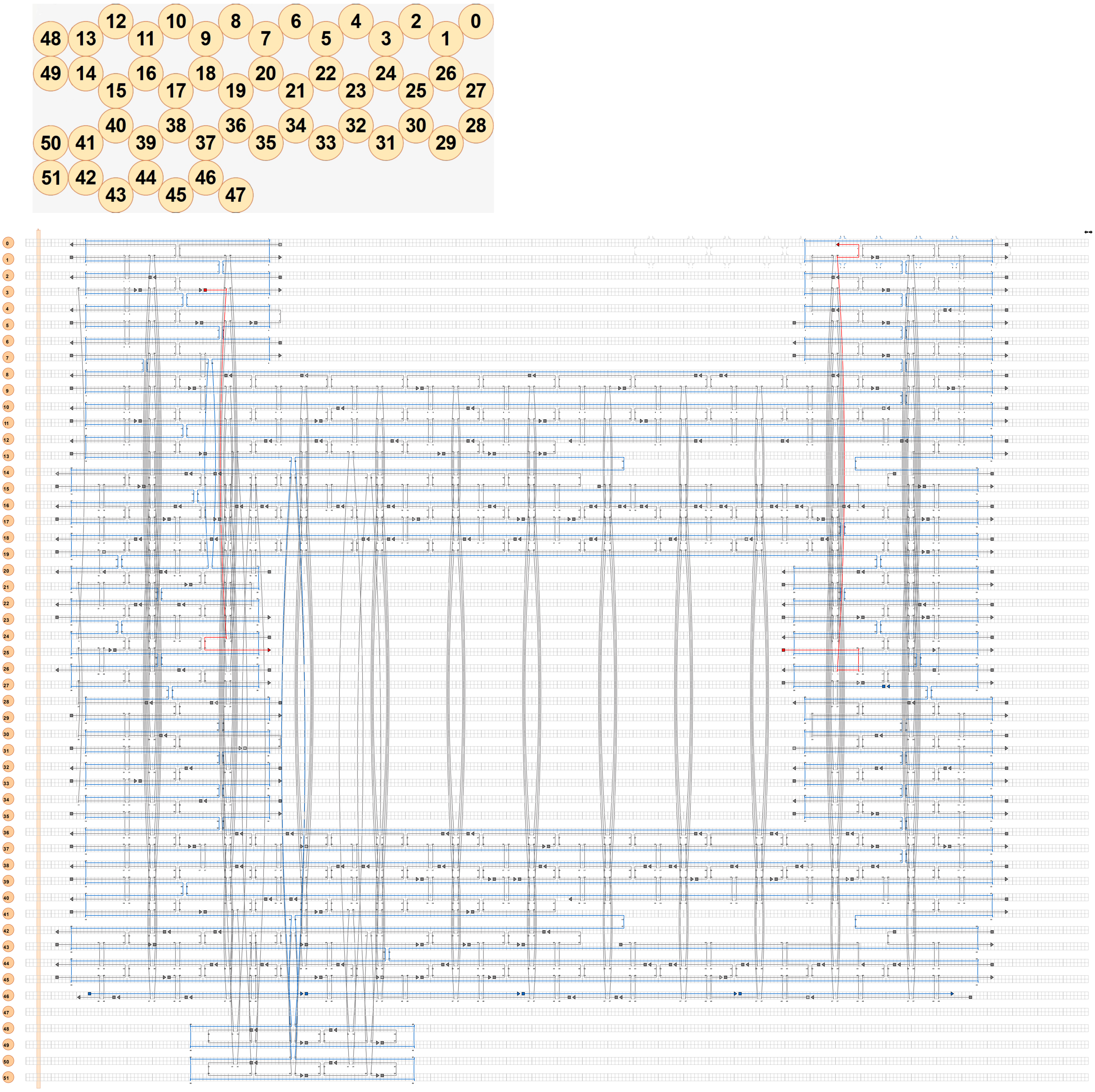

**Supplementary Figure 1. Design of the U-shape DNA origami frame.** Cross-section of the structure is shown on top of the strand diagram, both generated by caDNAno. Handles extend from the ends of the red staple strands.

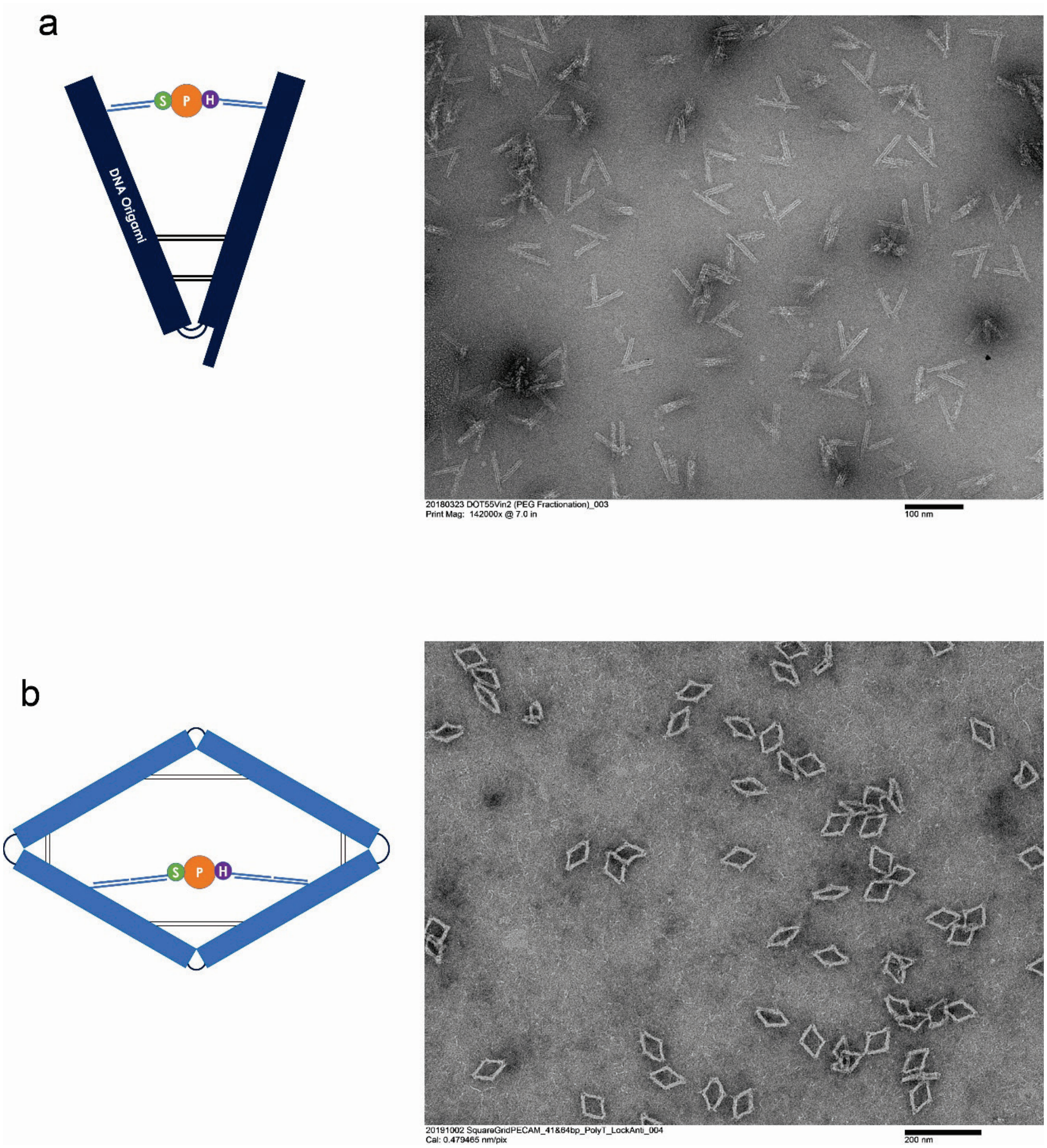


**Supplementary Figure 2. V- and rhombus-shape DNA origami frames. a)** Schematic and representative TEM image of V-shape frame for protein suspension. **b)** Schematic and representative TEM image of rhombus-shape frame for proteins suspension. In both schematics, blue rectangles represent multi-helical DNA origami arms and S-P-H represents the protein suspended in the DNA origami frame via double-stranded handles. Considerable structural deformation and heterogeneity can be seen from the negative-stain TEM images of protein-free DNA frames.

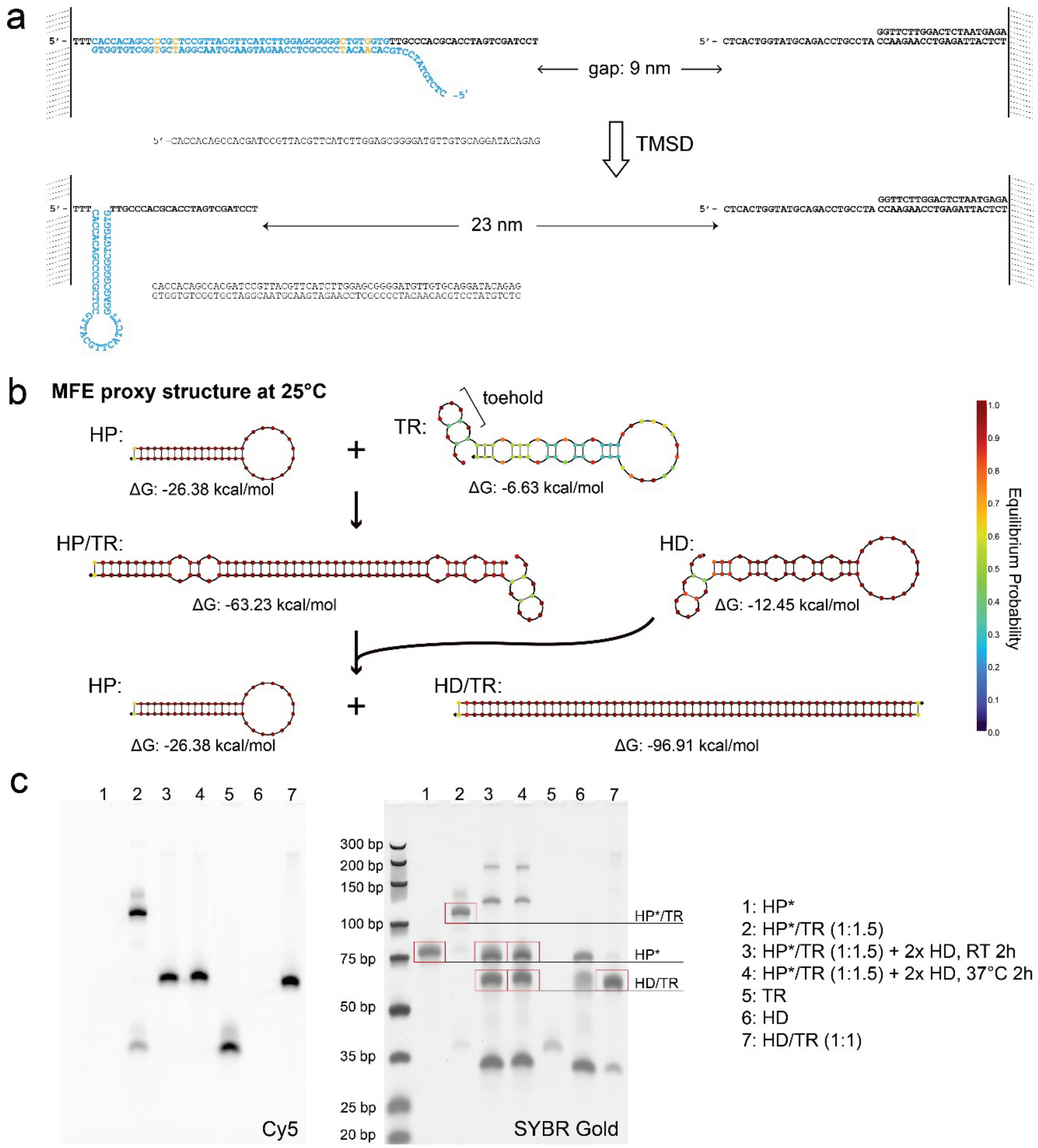


**Supplementary Figure 3. Design and validation of the Left DNA Spring. a)** Schematics showing sequences of a reconfigurable handle (left, i.e., DNA spring) and a fixed handle (right) in a DNA origami device. The reconfigurable handle is converted from extended to folded state by TMSD. **b)** Schematics showing the formation of a tension-loaded (i.e., extended) spring and its activation. HP: hairpin strand with a 17-bp stem (76.5% GC content) and a 14-nt loop; TR: target strand that binds to HP to form tension-loaded spring; HD: displacement strand that triggers HP folding. Minimum free energy (MFE) structures are predicted by NUPACK (25°C, 100 mM $Na^+$ and 12 mM $Mg^{2+}$). Base pair mismatches are designed to lower energy barriers for HP/TR binding and TMSD. **c)** Validation of reactions shown in **b** by non-denaturing PAGE. TR is Cy5-labeled. HP*: A 105-nt staple strand carrying the HP sequence and BG-tether-binding sequence.

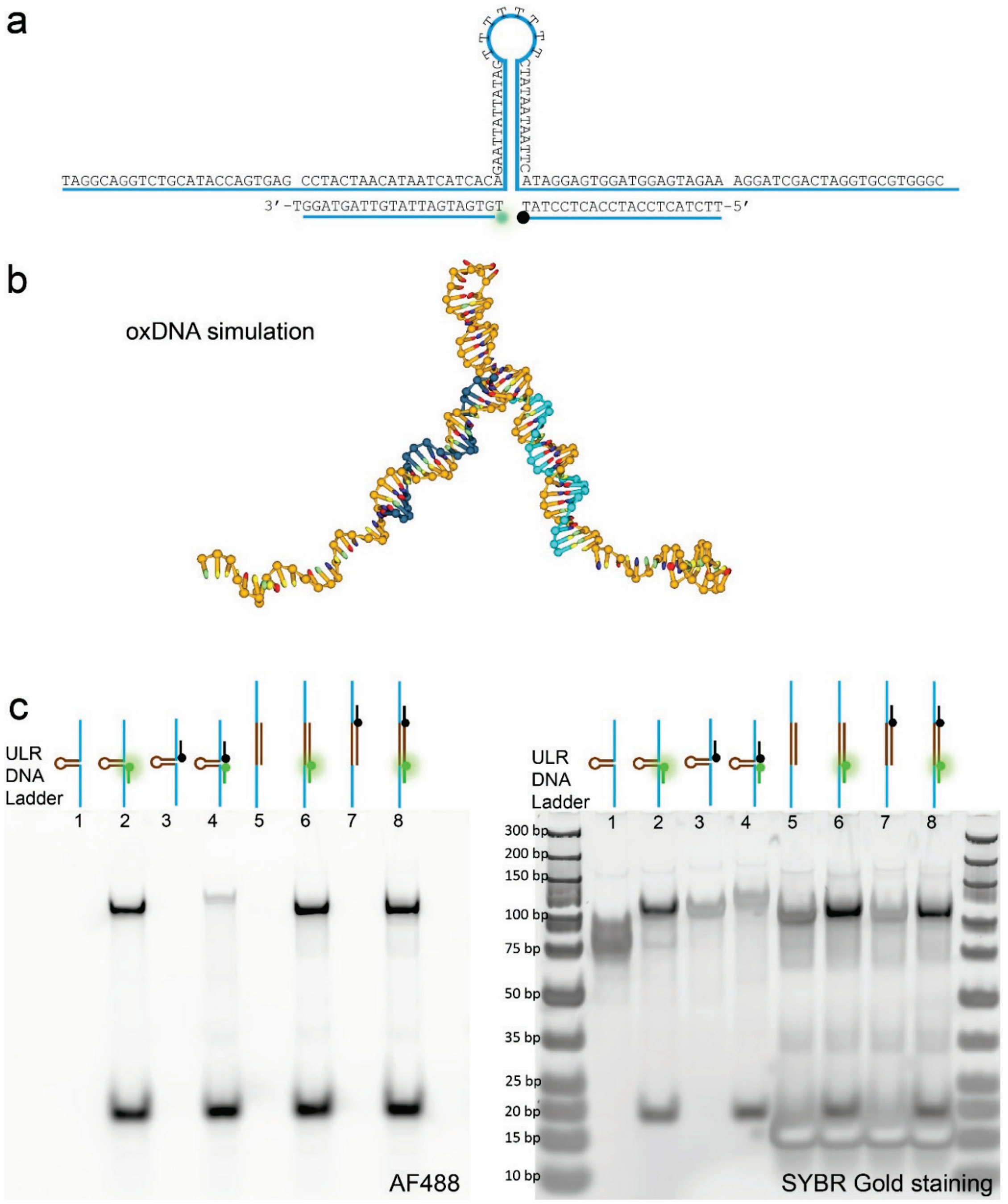


**Supplementary Figure 4. FRET-based DNAts. a)** DNAts (predicted unfolding force: ~8.3 pN) consisting of a 12-bp stem (17% GC content) and a $dT_8$ loop. Its maximum end-to-end distance in the folded state is ~16 nm, accounting for the lengths of BHQ1-DNA and AF488-DNA (21 nt each) and the stem diameter (2 nm). **b)** A snapshot of DNAts simulation by oxDNA. The relaxed 3-arm junction leads to a smaller end-to-end distance. **c)** Validation of DNAts assembly (lanes 1–4) by non-denaturing PAGE. As a control, an oligonucleotide with complementary stem-loop sequence was added in 10-fold excess during assembly (lanes 5–8).

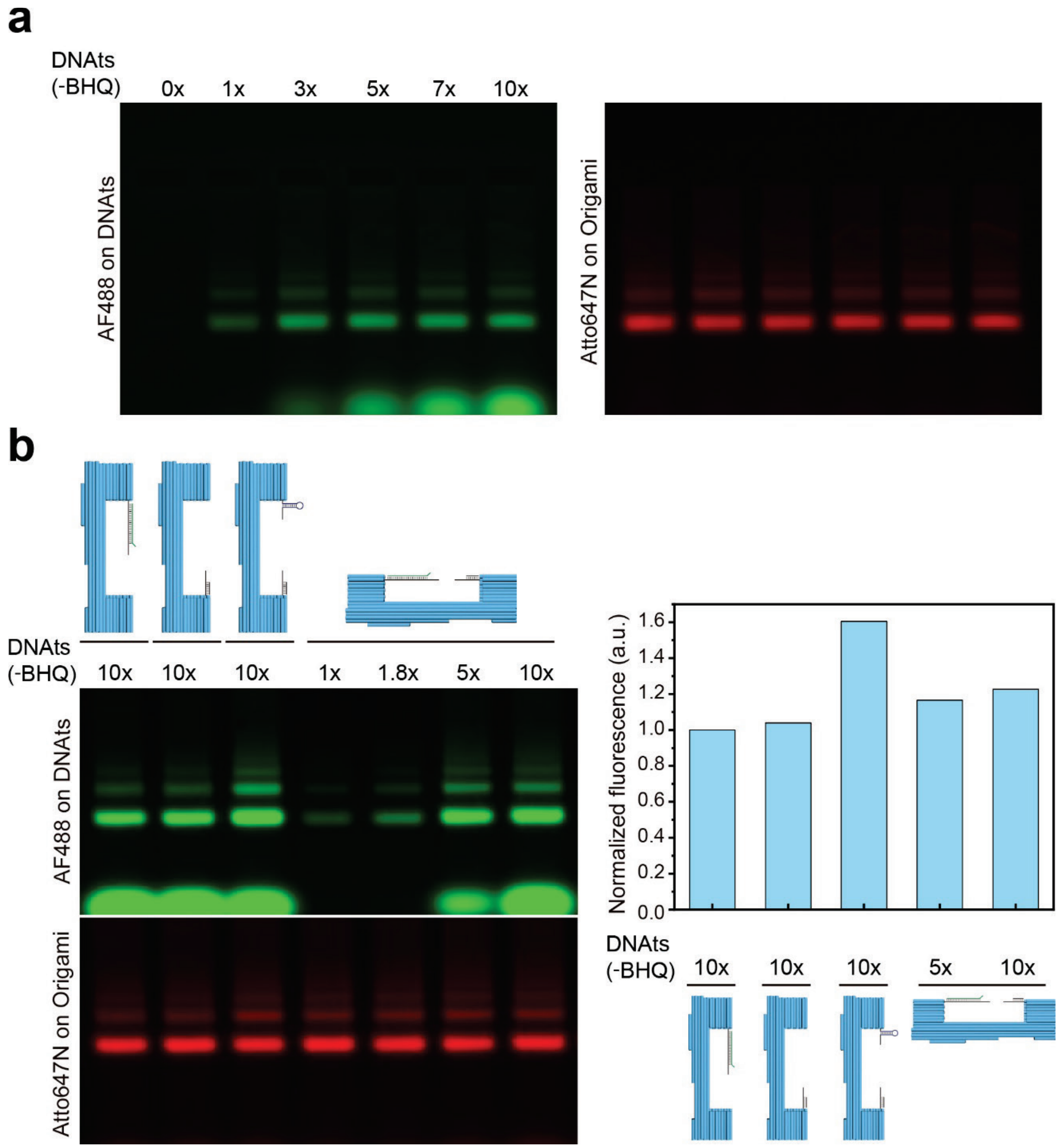


**Supplementary Figure 5. Loading DNAts into DNA origami device. a)** Loading efficiencies of DNAts into DNA origami device assessed by agarose gel electrophoresis at increasing DNAts:origami molar ratios. **b)** Loading efficiencies of DNAts into DNA origami devices with various handle configurations (see schematics on top of the gel images) assessed by agarose gel electrophoresis. DNAts is labeled with AF488 but without BHQ1; DNA origami device is labeled with Atto647N. Comparing the AF488/Atto647N ratios of the DNA device bands shows ~16% higher AF488 signal on DNA devices with two extended handles than those with only one handle, which we attribute to ~16% of the devices erroneously capturing two DNAts in its cavity.

**a**

50% GC DNAts: GAGTTCGTGTAG TTTTTTTT CTACACGAACTC; $F_{1/2} \approx 11.2$ pN

17% GC DNAts: GAATTATTATAG TTTTTTTT CTATAATAATTC; $F_{1/2} \approx 8.3$ pN

17% GC-1mismatch DNAts: GAATTAATATAG TTTTTTTT CTATAATAATTC; $F_{1/2} \approx 6.1$ pN

**b**

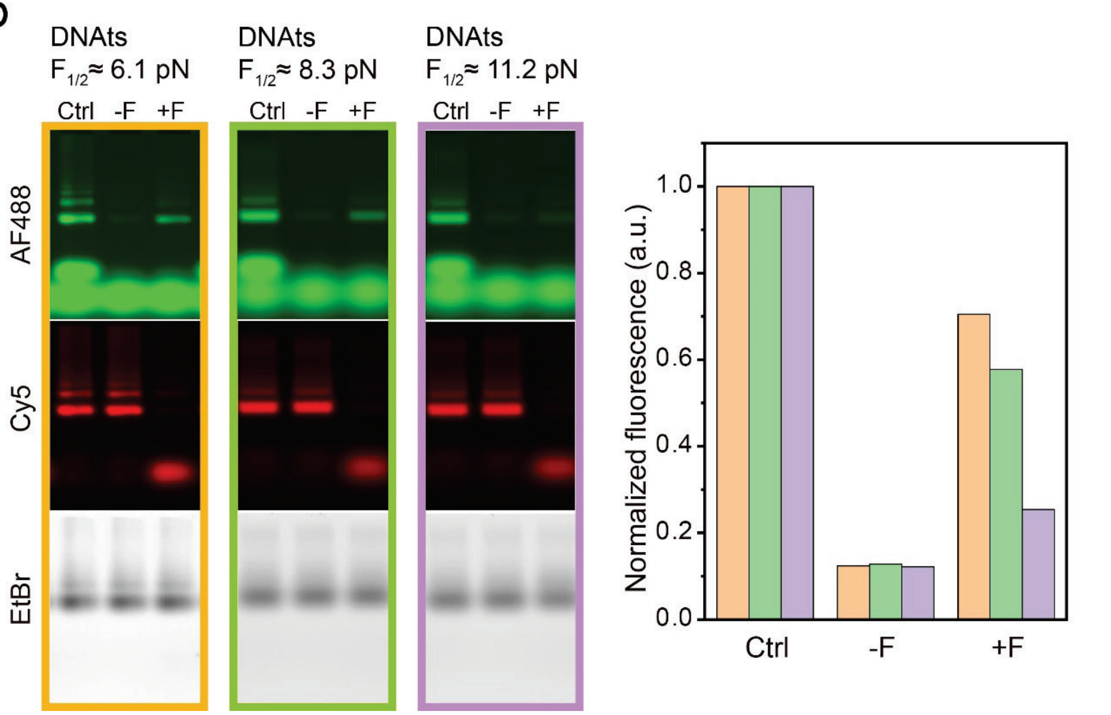


**Supplementary Figure 6. Unzipping DNAts with different unfolding forces. a)** DNAts designed with different GC contents and/or mismatch to tune their unfolding forces ($F_{1/2}$). **b)** Left: images of agarose gel (upper: AF488 channel, middle: Cy5 channel, lower: EtBr channel) in which the DNA devices loaded with DNAts missing BHQ1 (Ctrl) and with complete DNAts before (-F) and after (+F) TMSD were electrophoresed. Right: quantification of normalized AF488 signal on the left gel. The data for unzipping 8.3 pN DNAts (as shown in Figure 1c) are shown here again for comparison.

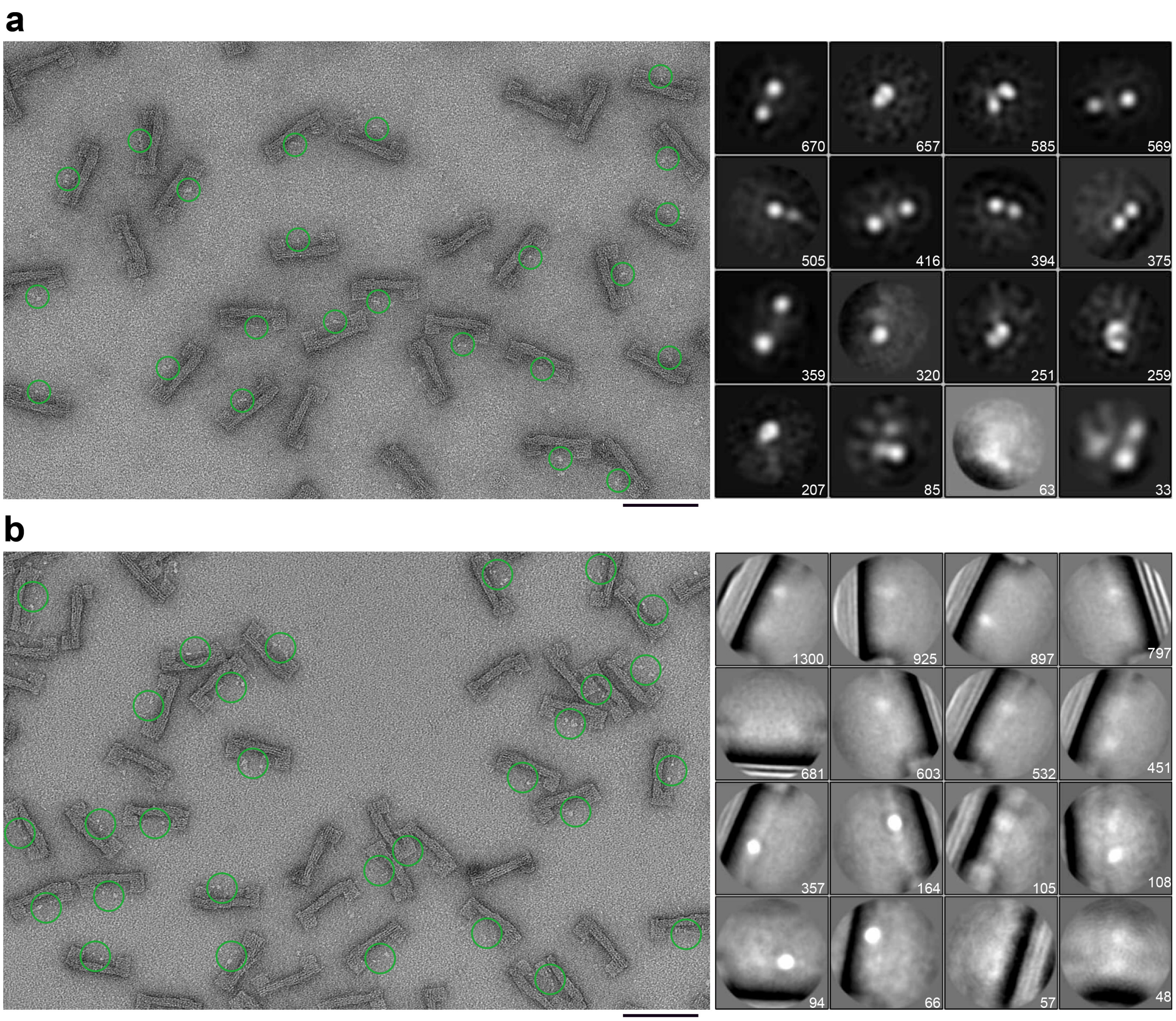


**Supplementary Figure 7. Additional TEM images of Origami-R1-R2 (a)** before (5,748 auto-picked particles) and **(b)** after (7,185 auto-picked particles) force application. Green circles in the zoomed-out images contain protein molecules selected for averaging. Averaged images are centered on the U-frame cavity where S-R1-R2-H is expected. Particle numbers are shown for each class. Selected averaged TEM images are shown in Figure 3. Scale bars: 100 nm. Box sizes: (a) 30 nm; (b) 50 nm.

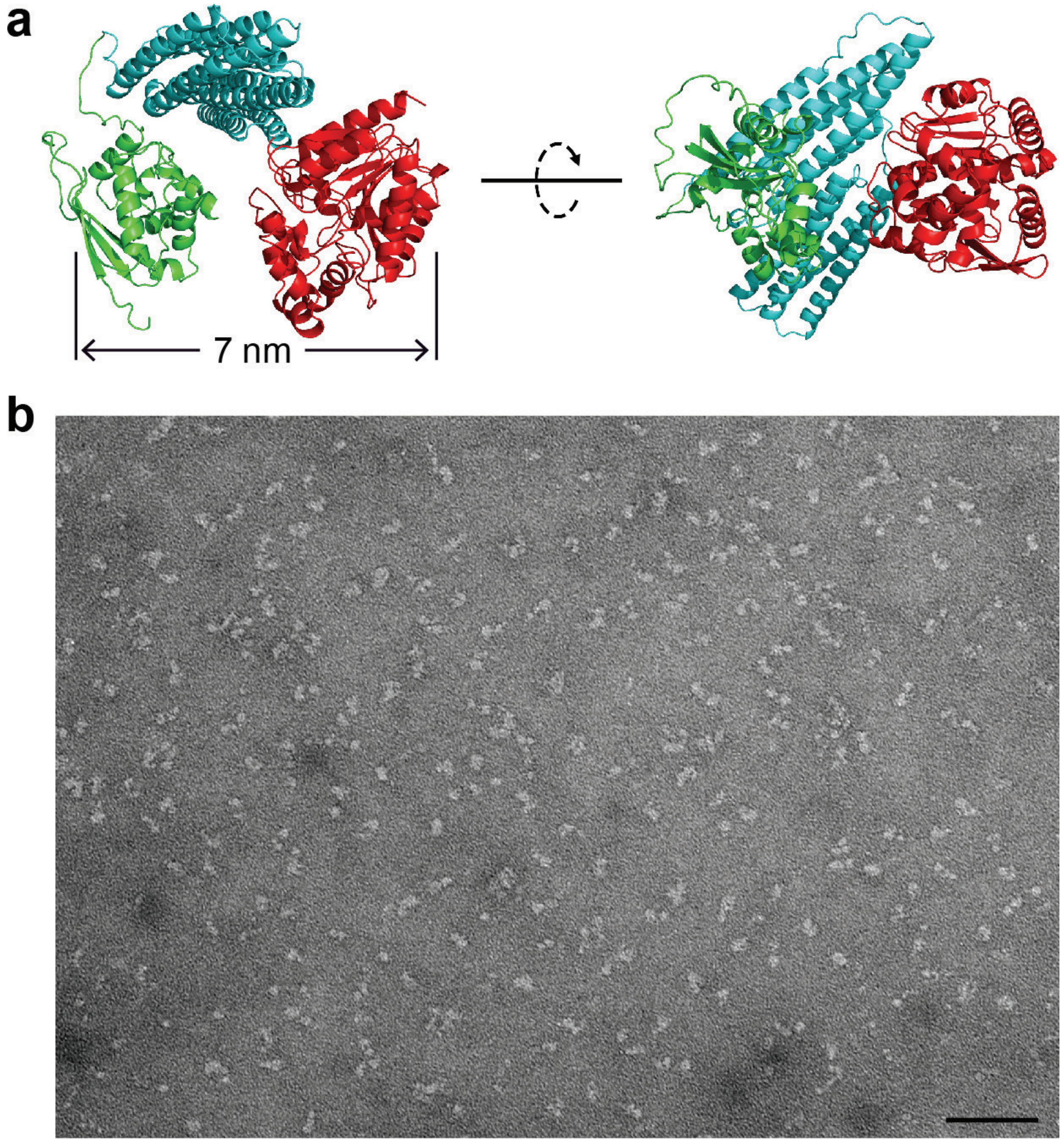


**Supplementary Figure 8. AlphaFold model and TEM characterization of S-R1-R2-H. a)** AlphaFold model of S-R1-R2-H showing three domains, SNAP-tag (green), R1-R2 (cyan), and Halo-tag (red), connected by short peptide linkers. **b)** A negative-stain TEM image of tether-conjugated S-R1-R2-H. Scale bar: 50 nm.

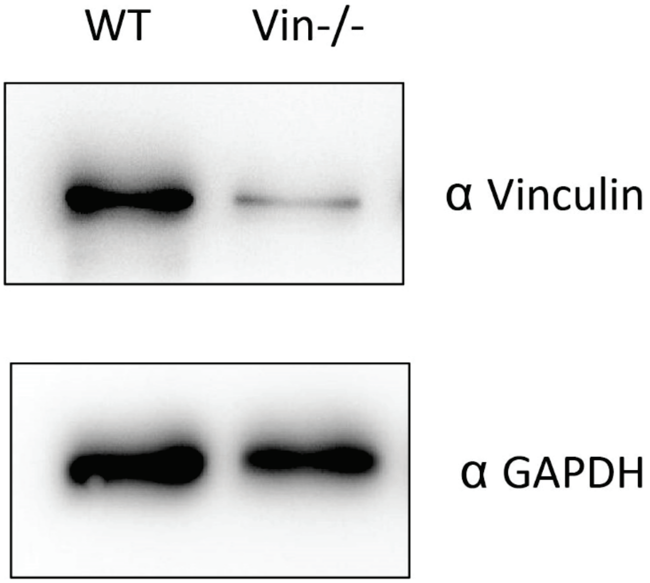


**Supplementary Figure 9. Vinculin knockdown efficiency.** Normalizing the vinculin band intensities by the GAPDH band intensities on a Western blot of wild-type (WT) and vinculin-depleted (Vin-/-) cell lysates shows a knockdown efficiency of ~87%**.**

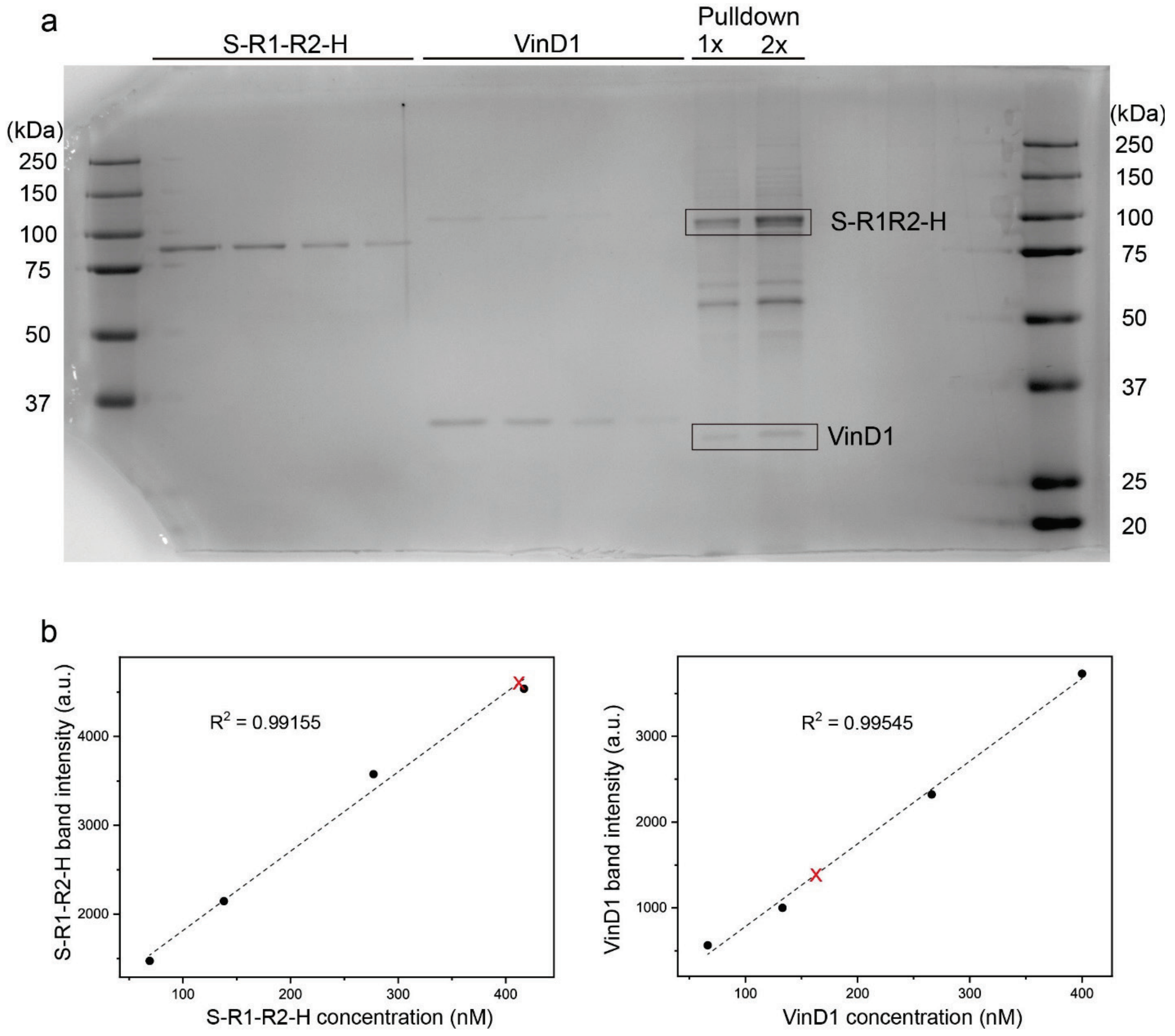


**Supplementary Figure 10. Quantitative analysis of S-R1-R2-H and VinD1 binding stoichiometry. a)** SDS-PAGE of purified S-R1-R2-H (unconjugated, 87.3 kDa) and VinD1-FLAG (expressed in *E. coli*, 33.5 kDa) loaded with a range of known concentrations, as well as the mixture of a pulldown experiment (6 nm gap, force applied using a 12.8 pN spring) loaded in two lanes with 2-fold concentration difference. **b)** Determining concentrations of S-R1-R2-H (left) and VinD1-FLAG (right) in the pulldown sample. Band intensities of the purified proteins (black dots, measured from **a**) were used to construct calibration lines by linear regression. The intensities of the corresponding protein bands in the pulldown sample (red crosses, measured from the 1× pulldown lane in **a**) were then used to determine the concentrations of S-R1-R2-H released from DNA devices and the pulled-down VinD1-FLAG (expressed in 293TX cells, 30.9 kDa). The resulting VinD1:R1R2 molar ratio is 163 nM/412 nM≈0.40.

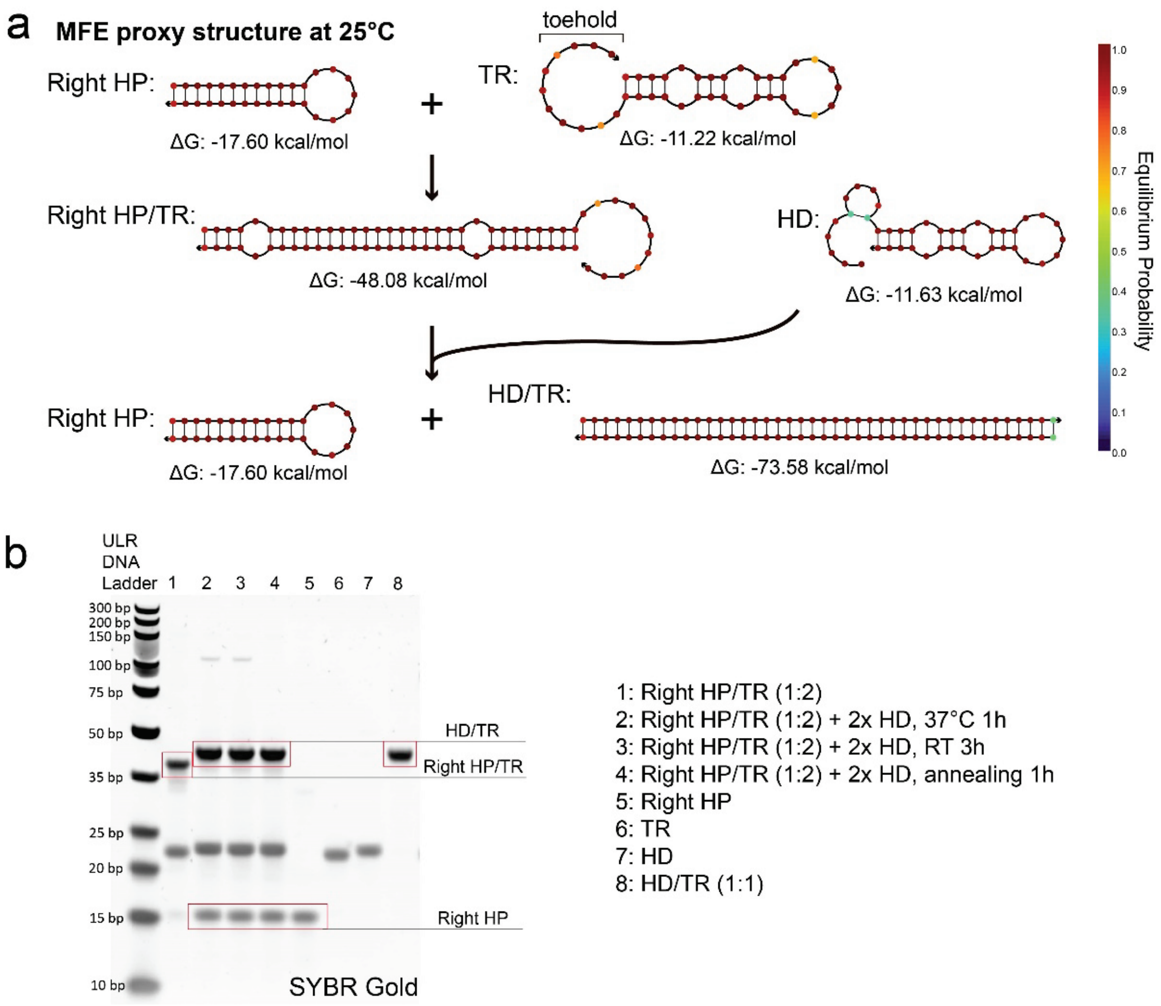


**Supplementary Figure 11. Design and validation of the Right DNA spring. a)** Schematics showing the formation of a tension-loaded (i.e., extended) right spring and its activation. HP: hairpin strand with a 12-bp stem (75% GC content) and a 7-nt loop; TR: target strand that binds to HP to form tension-loaded spring; HD: displacement strand that triggers HP folding. Minimum free energy (MFE) structures are predicted by NUPACK (25°C, 100 mM $Na^+$ and 12 mM $Mg^{2+}$). Base pair mismatches are designed to lower energy barriers for HP/TR binding and TMSD. **b)** Validation of reactions shown in **a** by non-denaturing PAGE.

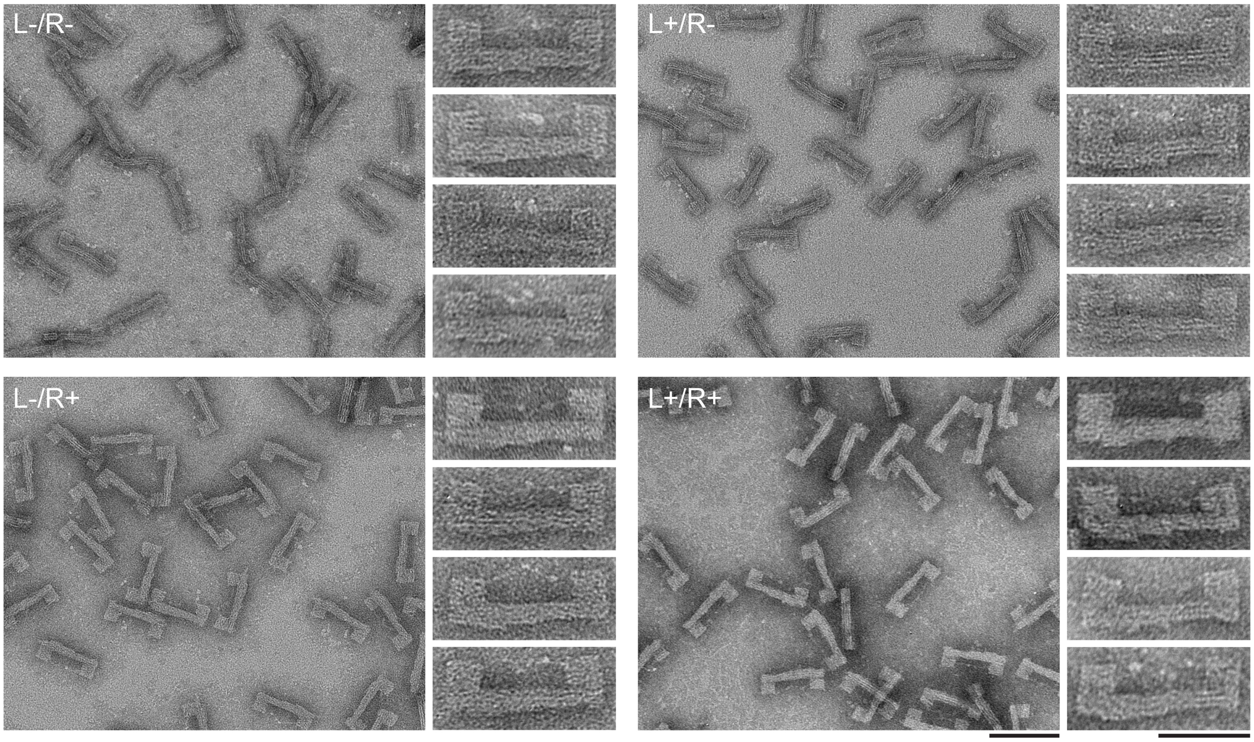


**Supplementary Figure 12. TEM images of S-R1-R2-H in the dual-spring DNA device.** The four groups of images show proteins in the relaxed (L-/R-) and various extended conditions (L+/R-, L-/R+, L+/R+). L/R denote left/right springs and -/+ denote loaded/activated state. Scale bars for zoomed-in and zoomed out images are 50 and 100 nm, respectively.

**Supplementary Table 1**. DNA oligonucleotides used for spring (spr) assembly and activation, DNAts assembly, and immobilizing DNA devices on streptavidin beads. Sequences are written from 5' to 3' end.

| | |
|---|---|
| For 12.8 pN spring (strong HP): | |
| Left77%GC Spr+7nm | GAATTCGCAGATGCGTGTTCAGCAAATCGT TTT CACCACAGCCCCGCTCC GTTACGTTCATCTT GGAGCGGGGCTGTGGTG TT ACTTCTGACGCCACGTCACTT GCCCACGCACCTAGTCGATCCT |
| Left+7nm_complement | AAGTGACGTGGCGTCAGAAGT |
| Left77%GC Spr | GAATTCGCAGATGCGTGTTCAGCAAATCGT TTT CACCACAGCCCCGCTCC GTTACGTTCATCTT GGAGCGGGGCTGTGGTG TT GCCCACGCACCTAGTCGATCCT |
| Left77%GC Spr-TR | CTCTGTATCCTG CACAACATCCCCGCTCC AAGATGAACGTAAC GGATCGTGGCTGTGGTG |
| Left77%GC Spr-TR_Cy5 | /5Cy5/CTCTGTATCCTG CACAACATCCCCGCTCC AAGATGAACGTAAC GGATCGTGGCTGTGGTG |
| Left77%GC Spr-HD | CACCACAGCCACGATCC GTTACGTTCATCTT GGAGCGGGGATGTTGTG CAGGATACAGAG |
| For 8.8 pN spring (weak HP): | |
| Left18%GC Spr | GAATTCGCAGATGCGTGTTCAGCAAATCGT TTT CTTTTAATTATTAGTTC GTTACGTTCATCTT GAACTAATAATTAAAAG TT GCCCACGCACCTAGTCGATCCT |
| Left18%GC Spr-TR | CTCTGTATCCTG CTTGTAAGTATTAGTTC AAGATGAACGTAAC GAAGTAACAATTAAAAG |
| Left18%GC Spr-HD | CTTTTAATTGTTACTTC GTTACGTTCATCTT GAACTAATACTTACAAG CAGGATACAGAG |
| For unstructured spring: | |
| Left_unstruct_Spr | GAATTCGCAGATGCGTGTTCAGCAAATCGT TTT TCCTTACTGCTTCTTGAGTTCACTGTTATCTTCATCGCGCTTCATACC TT GCCCACGCACCTAGTCGATCCT |
| Left_unstruct_Spr-TR | TGTGACCTAT GGTATGAAGTGCGATGAAGATAACAGTGAACTCAAGATGCAGTAAGGA |
| Left_unstruct_Spr-HD | TCCTTACTGCATCTTGAGTTCACTGTTATCTTCATCGCACTTCATACC ATAGGTCACA |
| For fixed handles: | |
| Right+0nm | CTCACTGGTATGCAGACCTGCCTA GCTGGCGAAAGGGGGATGTGTTTTCCAAGCCTTTGACCCT |
| Right+7nm | CTCACTGGTATGCAGACCTGCCTA CCAAGAACCTGAGATTACTCT GCTGGCGAAAGGGGGATGTGTTTTCCAAGCCTTTGACCCT |
| Right+7nm_complement | AGAGTAATCTCAGGTTCTTGG |
| Rright+10nm | CTCACTGGTATGCAGACCTGCCTA TAGATCGGACCAAGAACCTGAGATTACTCT GCTGGCGAAAGGGGGATGTGTTTTCCAAGCCTTTGACCCT |
| Right+10nm_complement | AGAGTAATCTCAGGTTCTTGGTCCGATCTA |
| For right spring: | |
| Right75%GC Spr | CTCACTGGTATGCAGACCTGCC CGCTACGGTCGC CTGAGAT GCGACCGTAGCG T GCTGGCGAAAGGGGGATGTGTTTTCCAAGCCTTTGACCCT |
| Right75%GC-HP | CGCTACGGTCGC CTGAGAT GCGACCGTAGCG |
| Right75%GC Spr-TR | CGCTTCGGTCGC ATCTCAG GCGTCCGTAGCG GCTGTCGGAAT |
| Right75%GC Spr-HD | ATTCCGACAGC CGCTACGGACGC CTGAGAT GCGACCGAAGCG |
| For DNAts: | |

| | |
|---|---|
| DNAts 50%GC | TAGGCAGGTCTGCATACCAGTGAG CCTACTAACATAATCATCACA GAGTTCGTGTAG TTTTTTTT CTACACGAACTC ATAGGAGTGGATGGAGTAGAA AGGATCGACTAGGTGCGTGGGC |
| DNAts 17%GC | TAGGCAGGTCTGCATACCAGTGAG CCTACTAACATAATCATCACA GAATTATTATAG TTTTTTTT CTATAATAATTC ATAGGAGTGGATGGAGTAGAA AGGATCGACTAGGTGCGTGGGC |
| DNAts 17%GC S12T8-1mis | TAGGCAGGTCTGCATACCAGTGAG CCTACTAACATAATCATCACA GAATTAATATAG TTTTTTTT CTATAATAATTC ATAGGAGTGGATGGAGTAGAA AGGATCGACTAGGTGCGTGGGC |
| DNAts_BHQ1 | TTCTACTCCATCCACTCCTAT/3BHQ_1/ |
| DNAts_Ctrl | TTCTACTCCATCCACTCCTAT |
| DNAts_AF488 | /5Alex488N/TGTGATGATTATGTTAGTAGGT |
| DNAts 17%GC_complement | GAATTATTATAGAAAAAAAACTATAATAATTC |
| DNAts_50%GC_complement | GAGTTCGTGTAGAAAAAAAACTACACGAACTC |
| For DNA device pulldown: | |
| Biotin-anchor staple1 | GCG GTC TCA ATC AGG CAT GGA GTA C ACGAACCACAGTGCCACGTTTT |
| Biotin-anchor staple2 | GCG GTC TCA ATC AGG CAT GGA GTA C CTGAGAGCCTATCAAAATTTTT |
| Biotin-anchor staple3 | GCG GTC TCA ATC AGG CAT GGA GTA C TTTTCTTTTTCAAATATCCTTATCATTCTTTT |
| Biotin-anchor staple4 | GCG GTC TCA ATC AGG CAT GGA GTA C TTTTCAAGAACGGGTACAAGCCGTTTTTTTTT |
| Biotin-labeled strand | GTA CTC CAT GCC TGA TTG AGA CCG C TTT/3Bio/ |
| DNA tethers: | |
| BG-tether | /5AmMC6/AGGATCGACTAGGTGCGTGGGC |
| CL-tether | TAGGCAGGTCTGCATACCAGTGAG /3AmMO/ |
